# Reactive Astrocytes Drive Extracellular Acidification to Mediate α-Synuclein Neurodegeneration

**DOI:** 10.64898/2026.01.06.697893

**Authors:** Jae-Jin Song, Hyejin Park, YuRee Choi, Taekyung Ryu, Jisu Shin, Seo-Hyun Kim, Albert Park, Justin Wang, Devanik Biswas, Shih-Ching Chou, Shinwon Ha, Yura Jang, Yu Shin, Hao Wen Chen, Ingie Hong, John Wemmie, Per Svenningsson, Juan Troncoso, Jiadi Xu, Chan Hyun Na, Valina L. Dawson, Ted M. Dawson, Tae-In Kam

## Abstract

Astrocytes are increasingly recognized as key players in neurodegeneration^1–3^, yet the molecular mechanisms by which they drive disease remain elusive. Here, we uncover a fundamental pathway in which reactive astrocytes fuel neurodegeneration in α-synucleinopathies—including Dementia with Lewy bodies and Parkinson’s disease dementia—by acidifying the brain’s extracellular environment. We demonstrate that both human patient tissue and a gut-to-brain α-synuclein mouse model exhibit accumulation of reactive astrocytes and extracellular acidosis. Mechanistically, we show that astrocytic lysosomal exocytosis releases acidic contents, driving a drop in pH that activates neuronal acid-sensing ion channel 1a (ASIC1a), resulting in neuronal loss and behavioral decline. Blocking this pathway—either by inhibiting astrocytic lysosomal exocytosis or genetically or pharmacologically targeting neuronal ASIC1a—mitigates pathology and rescues neurodegenerative phenotypes in vivo. These findings provide a conceptual advance by establishing a mechanistic link between glial inflammation, acid-base homeostasis, and neuronal vulnerability, and suggest that targeting astrocyte-driven acidification or ASIC1a signaling could offer new avenues for disease modification in α-synucleinopathies.

## Main

Synucleinopathies are a group of neurodegenerative disorders defined by the abnormal accumulation of the protein α-synuclein in the brain ^4,5^. Dementia with Lewy bodies (DLB) and Parkinson’s disease dementia (PDD) are clinically distinct forms of synucleinopathies collectively referred to as Lewy body dementias ^6–8^. Both conditions are characterized by progressive cognitive and motor decline, yet they differ in the timing and pattern of symptom onset and disease progression ^6,7^. Despite these clinical distinctions, they share the pathological hallmark of widespread α-synuclein aggregation and neuronal loss, pointing to shared mechanisms of neurodegeneration that remain incompletely understood.

Astrocytes are the most abundant glial cells in the central nervous system and play essential roles in supporting neuronal function, maintaining homeostasis, and modulating synaptic activity ^9–11^. In neurodegenerative conditions, astrocytes undergo morphological and transcriptional changes to become reactive in a heterogeneous and context-dependent manner ^1,12^. Pathological α-synuclein or oligomeric amyloid-β triggers microglial activation, which in turn drives the formation of activated neurotoxic reactive (NTR) astrocytes that damage neurons in models of Parkinson’s disease and Alzheimer’s disease ^2,13^. Inhibiting microglial activation suppresses this harmful astrocyte state and mitigates neurodegeneration both in vitro and in vivo ^1,2,13^. NTR astrocytes are emerging as key modulators of neurodegenerative disease pathophysiology, yet the precise cellular and molecular mechanisms by which they directly contribute to neurodegeneration remain unclear.

Here, we uncover a mechanism by which NTR astrocytes promote α-synuclein-induced neurodegeneration. We show that NTR astrocytes accumulate in postmortem brains in a gut-to-brain transmission mouse model of α-synucleinopathy and in patients with Lewy body dementias. Genetic ablation of microglia-derived cytokines required for NTR astrocyte induction - interleukin-1α (IL-1α), tumor necrosis factor α (TNFα), and C1q -markedly reduces α-synuclein pathology, neuronal loss, and behavioral deficits in the gut-to-brain model of pathologic α-synuclein-induced degeneration, supporting a critical role for microglia-driven NTR astrocyte generation in disease pathogenesis. Proteomic profiling of the NTR astrocyte secretome identified lysosomal proteins as a major component of the NTR astrocyte secretome. Mechanistically, the extracellular release of lysosomal contents by NTR astrocytes leads to extracellular acidification. This acidic microenvironment activates neuronal acid-sensing ion channels 1 (ASIC1), triggering neuronal cell death, synaptic degeneration, and α-synuclein aggregation. Cerebrospinal fluid from patients with Lewy body dementias and brain tissue from the gut-to-brain model of pathologic α-synuclein showed significant acidosis, supporting the link between NTR astrocyte mediated extracellular acidification and disease pathology. Pharmacological inhibition or genetic deletion of ASIC1a mitigates NTR astrocyte-mediated neurotoxicity, reduces synaptic loss, and alleviates motor and cognitive impairments in gut-to-brain model of pathologic α-synuclein induced degeneration. These findings highlight NTR astrocytes as non-neuronal drivers of neurodegeneration by acidifying the extracellular environment and suggest that targeting NTR astrocyte lysosomal exocytosis or neuronal ASIC1a activity may offer a therapeutic avenue for α-synucleinopathies and potentially other neurodegenerative diseases in which NTR astrocytes are implicated.

## Results

### NTR astrocytes are present in the Lewy body dementia brains and the gut-brain transmission model of pathologic α-synuclein neurodegeneration

NTR astrocytes are characterized by C3 immunoreactivity of glial fibrillary acidic protein (GFAP)-positive cells and by distinct transcriptional signatures associated with neurodegeneration in several human degenerative brain disorders ^1^. To determine whether NTR astrocytes are present in Lewy body dementia, we examined brain tissue from patients with DLB and PDD (Extended Data Fig. 1a and Table S1). Immunohistochemistry confirmed a significant accumulation of C3⁺ and GFAP⁺ astrocytes in the midbrain (Extended Data Fig. 1b-d) and in the hippocampus (Extended Data Fig. 1e–g) in both disorders. Transcriptomic analysis of astrocytes enriched by immunopanning revealed an upregulation of NTR astrocyte-associated gene expression profiles consistent with the presence of NTR astrocytes in DLB and PDD (Extended Data Fig. 1h, i).

To model this process in vivo, we used a gut-to-brain pathologic α-synuclein transmission mouse model, in which α-synuclein preformed fibrils (α-syn PFFs) were injected into the pylorus and duodenum, triggering conversion of endogenous α-synuclein to pathogenic α-synuclein, which propagates via the vagus nerve to the brain ^14,15^. This model recapitulates the full features of α-synucleinopathy induced degeneration, including α-syn aggregation, dopaminergic neurodegeneration, and motor and cognitive impairments. We assessed the induction of NTR astrocytes in the ventral midbrain and hippocampus at 2, 6, and 10 months following α-syn PFF injection (Extended Data Fig. 1a). C3⁺ and GFAP⁺ NTR astrocytes were evident as early as 2 months post-injection, preceding detectable neurodegeneration or behavioral deficits (Extended Data Fig. 1j–o). The abundance of C3⁺ and GFAP⁺ cells increased progressively at 6 and 10 months. Expression of NTR astrocyte-associated transcripts were also significantly elevated from 2 months and progressively increased at 6 and 10 months (Extended Data Fig. 1p, q). These findings suggest that pathological α-syn induce a time-dependent expansion of NTR astrocytes that parallels the progression of α-syn pathology in this model ^14,15^, suggesting a temporal association between astrocytic reactivity and α-syn pathology.

### Genetic deletion of NTR astrocyte-inducing factors prevents NTR astrocyte formation and mitigates α-synucleinopathy-associated neurodegeneration

NTR astrocytes are induced by activated microglia through secretion of IL-1α, TNFα, and C1q—cytokines that are collectively necessary and sufficient to trigger this astrocyte phenotype ^1^. Microglia activated by α-syn PFFs or amyloid-β oligomers drive NTR astrocyte formation in models of PD and AD, respectively ^2,13,16^ . To investigate the causal role of NTR astrocyte inducing factors (IL-1α, TNFα, and C1q) in pathologic α-synuclein induced degeneration, mice deficient in *Il1a*, *Tnf* and *C1qa* (triple KO, tKO) were subjected to the gut α-syn PFF injection model (Fig. 1a). Eight months post-injection, wild type (WT) mice injected with α-syn PFFs exhibited significant motor deficits as determined by the pole test (Fig. 1b, Extended Data Fig. 2b), grip strength test (Fig. 1c, Extended Data Fig. 2c) and open-field test (Extended Data Fig. 2d-h), and cognitive impairments as determined by Y-maze (Fig. 1d, e) and Morris water maze test (Fig. 1f-h), whereas tKO mice showed markedly improved performance across all assays (Fig. 1b-h). Pathological α-syn accumulation, measured via phosphorylated α-syn (pSer129-α-syn) immunostaining, was robustly increased in multiple brain regions in WT mice following gut-PFF injection including the prefrontal cortex (PFC), hippocampus (HIP), locus coeruleus (LC), amygdala (AMG), substantia nigra pars compacta (SNc), and striatum (STR), but was significantly reduced in tKO mice (Fig. 1i, j). In addition, dopamine (DA) neuron loss in the SNc, assessed via tyrosine hydroxylase (TH) and Nissl immunostaining and stereological counting, was evident in WT mice, but was markedly attenuated in tKO animals (Fig. 1k-m). Striatal TH immunoreactivity (Fig. 1n, o) and DA levels (Fig. 1p) also revealed significant DA neuronal loss in littermate control mice after gut-PFF injection. In contrast, reduction of striatal TH and DA levels was prevented in tKO mice (Fig. 1n-p).

**Figure 1.**
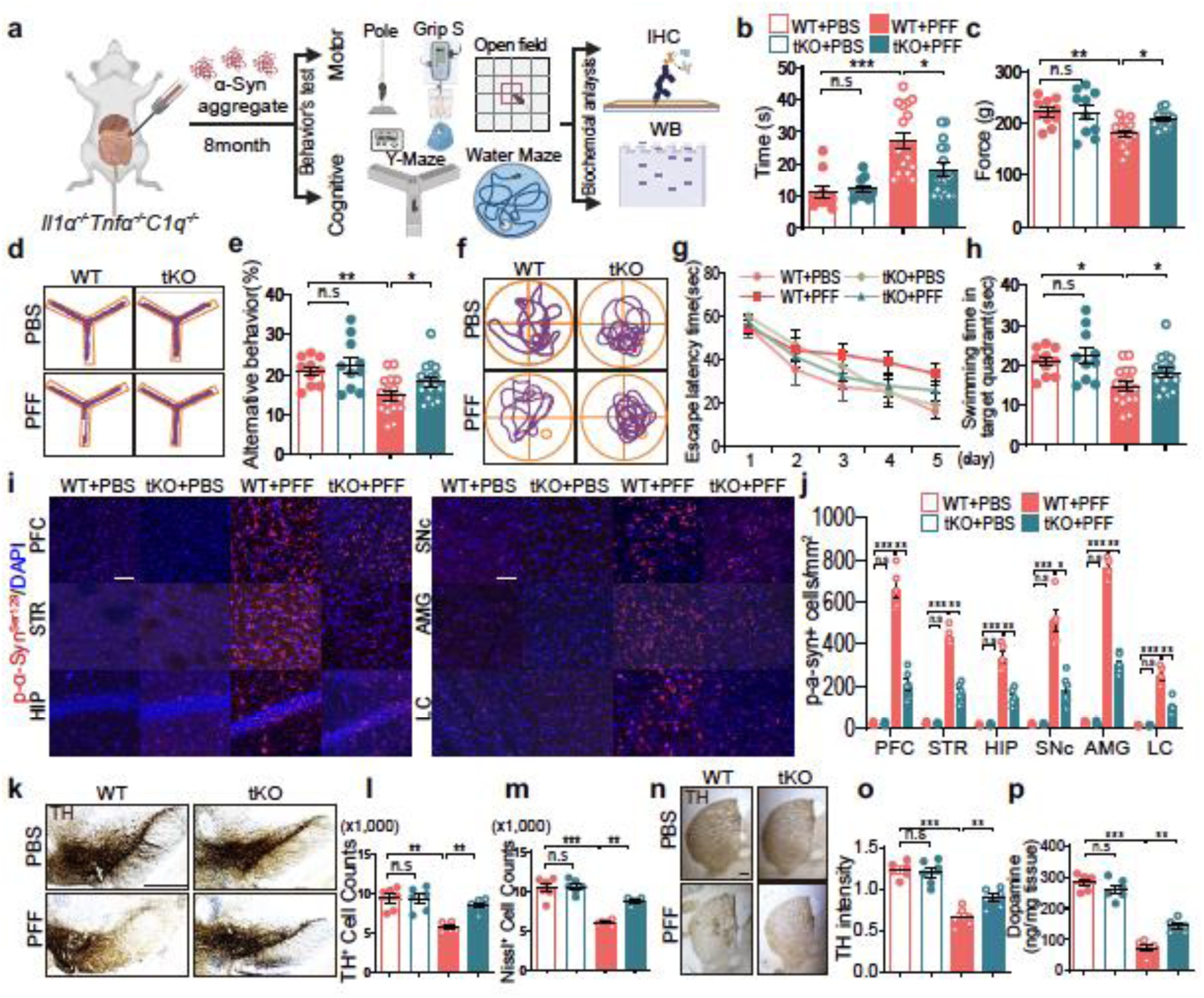
Lewy body dementia-like pathology in the gut-to-brain α -syn PFF model is rescued by genetic depletion of IL1α, TNFα and C1q. **(a)** Schematic diagram of the gastrointestinal injection of α-syn PFF or PBS in WT and tKO (*Il1a*, *Tnfa* and *C1qa KO*) mice. The figure was created with BioRender. **(b, c)** Motor function was measured by (b) pole test and (c) grip strength test 8 months after PBS and α-syn PFF injection. Data are the means ± s.e.m. Two-way ANOVA followed by Bonferroni’s post hoc test (n=10-16 mice for WT, n=13-16 mice for tKO mice). **(d-h)** Cognitive function was measured by (d, e) Y-maze test and (f-h) Morris water maze test 8 months after PBS and α-syn PFF injection. (d) Representative movement paths of mice from each group in the Y-maze test. (e) Percentage of alternative behavior in the Y-maze test. (f) Representative swimming paths of mice from each group in the Morris water maze test on the probe trial day 5. (g) Result of mice on the Escape latency time and (h) probe trial session in the Morris water maze test. Data are the means ± s.e.m. Two-way ANOVA followed by Bonferroni’s post hoc test (n=10-16 mice for WT, n=13-16 mice for tKO mice). **(i)** Distribution of pSer129-α-syn accumulation in the brains of PBS or α-syn PFF-injected mice. pSer129-α-syn immunohistochemistry from the dorsal motor nucleus of the vagus to the olfactory bulb of α-syn PFF gastrointestinal-injected mice sacrificed at 10 months post-injection (Ctx, cortex; DMV, dorsal motor nucleus of the vagus; HIP, hippocampus; LC, locus coeruleus; PFC, prefrontal cortex; SNc, substantia nigra pars compacta; STR, striatum). Scale bar, 50 μm. **(j)** Quantification of pSer129-α-syn immunoreactivity. Data are the means ± s.e.m. Two-way ANOVA followed by Bonferroni’s post hoc test (n=6). **(k)** Representative photomicrographs from coronal mesencephalon sections containing TH-positive neurons in the SNc region. Scale bar, 400 μm. **(l, m)** Unbiased stereological counts of (l) TH and (m) TH and Nissl positive in the SNc region. Data are the means ± s.e.m. Two-way ANOVA followed by Bonferroni’s post hoc test (n=6). **(n, o)** Representative photomicrographs from striatum region of WT and tKO mice 10 months after gut injection of α-syn PFF or PBS. Scale bar, 1mm. Quantification graphs of TH-intensity. Data are the means ± s.e.m. Two-way ANOVA followed by Bonferroni’s post hoc test (n=6). **(p)** DA concentration in the striatum of WT and tKO mice at 10 months after gut injection of α-syn PFF or PBS measured by ELISA. Data are the means ± s.e.m. Two-way ANOVA followed by Bonferroni’s post hoc test (n=6). *P < 0.05, **P < 0.01, ***P < 0.001,. n.s., not significant.

In WT mice subjected to gut-PFF injection, both pan-reactive and NTR astrocytes-related transcripts were strongly upregulated. However, this induction was significantly blunted in the SNc and hippocampus of tKO mice (Extended Data Fig. 3a, b). In contrast, transcripts specific to non-NTR phenotypes remained unchanged between genotypes, indicating a selective suppression of the NTR astrocytes (Extended Data Fig. 3a, b).

Immunohistochemistry revealed a substantial increase in GFAP^+^ and C3^+^ astrocytes in the SNc and hippocampus of WT mice injected with α-syn PFFs, which was notably suppressed in tKO mice (Extended Data Fig. 3c-h). Similar reductions in NTR astrocyte markers were also observed in other affected regions such as the prefrontal cortex and striatum (Extended Data Fig. 3i-n). Together, these findings demonstrate that the induction of NTR astrocytes via IL-1α, TNFα and C1q is critical for the development of α-syn pathology, DA neuronal loss, and the associated behavioral deficits.

### NTR astrocytes release lysosomal contents to the extracellular space

To explore how NTR astrocytes contribute to neurodegeneration, secretome profiling of astrocyte-conditioned media (ACM) derived from astrocytes treated with microglia-conditioned media (MCM) previously stimulated with α-syn PFFs, amyloid-β (Aβ) oligomers or lipopolysaccharide (LPS) known to induce NTR astrocytes was performed ^12,17^ (Fig. 2a).

**Figure 2.**
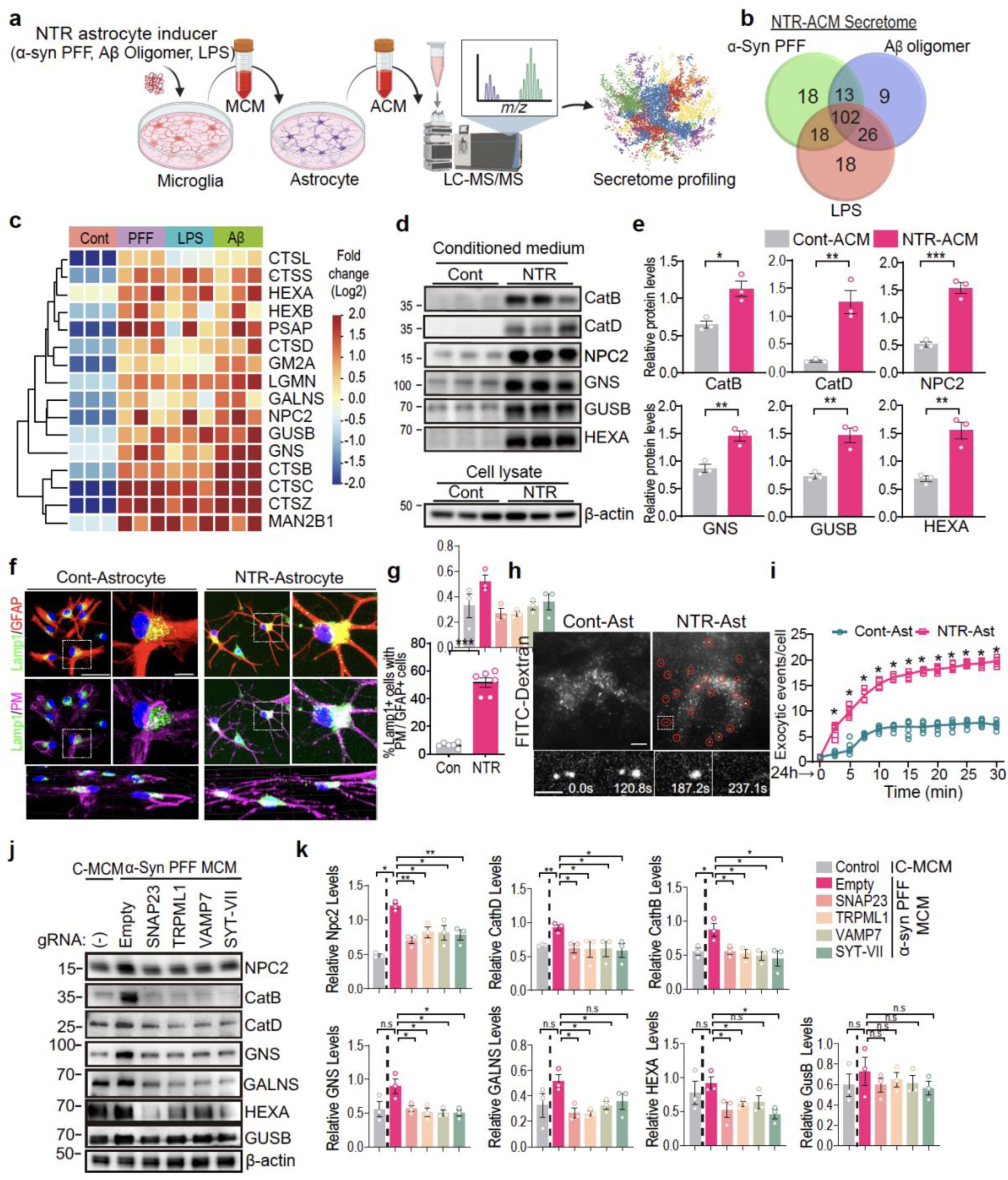
NTR astrocytes release lysosomal contents to extracellular space. **(a)** Schematic diagram of NTR-astrocytes secretome analysis using LC-MS/MS. The P values between the comparison groups were calculated using the student’s two-sample t-test. Proteins with q-values < 0.05 were considered differentially expressed (n=3). The figure was created with BioRender. **(b)** Venn diagram summarizing the secreted proteins from α-syn PFF (green), Aβ oligomer (blue) and LPS (red)-activated NTR astrocytes. **(c)** Lysosomal proteins detected from 102 overlapped proteins among α-syn PFF, Aβ oligomer and LPS-activated ACM. **(d, e)** Representative Immunoblots of Cathepsin B and D, NPC2, GNS, GUSB, HEXA, GALNS in ACM and β-actin in cell lysate. Quantification of secreted proteins levels normalized to β-actin Data are the means ± s.e.m. Unpaired two-tailed student *t* test (n=3). **(f, g)** Immunocytochemical analyses for the positioned lysosome, Lamp1(green) in astrocyte, GFAP(red) and Plasma membrane(purple). Scale bars, 50μm and 10μm for low and high magnification images. Quantification of Lamp1 positive cells out of plasma membrane. Data are the means ± s.e.m. Unpaired two-tailed student *t* test (n=6). **(h-i)** Representative TIRF images for the exocytic lysosome (FITC-Dextran) in NTR astrocytes. Scale bar, 10 μm and 1 μm for low and high magnification images. Quantification of exocytic event per cell with each time point in the control and NTR astrocytes. Data are the means ± s.e.m. Unpaired two-tailed student *t* test (n=6). **(j, k)** Representative immunoblots of Cathepsin B and D, NPC2, GNS, GUSB, HEXA, GALNS in primary mouse astrocytes infected with lentivirus containing CRISPR/Cas9 gRNAs targeting mock and lysosomal exocytosis-related genes. Quantification of the proteins level normalized to ponceau. Data are the means ± s.e.m. Two-way ANOVA followed by Bonferroni’s post hoc test (n=3). *p < 0.05; **p < 0.01; ***p < 0.001.

Approximately half of the secreted proteins (102 in total) from astrocytes stimulated by each inducer overlapped, suggesting the presence of shared neurotoxic factors released by NTR astrocytes (Fig. 2b, Extended Data Fig. 4a, b and Table S2). LC-MS/MS analysis of overlapping proteins revealed enrichment of gene ontologies related to immune response, system process and aging, and KEGG pathway associated with lysosome, phagosome, complement and coagulation cascades (Extended Data Fig. 4c). NTR astrocytes directly induced by the IL-1α, TNFα and C1q (ITC) cytokine cocktail secreted a similar set of proteins, indicating that the shared NTR astrocytes secretome is intrinsic to astrocyte activation and not derived from the microglia-conditioned media (Extended Data Fig. 4a, d).

Among 102 overlapped proteins, 16 lysosomal proteins were detected in each secretome of α-syn PFF-, amyloid-β oligomers-, or LPS-induced ACM (Fig. 2c). Significantly elevated secretion of cathepsins B and D, NPC2, and multiple lysosomal hydrolases were observed in the extracellular medium from α-syn PFF- or ITC cytokine-induced NTR astrocytes as assessed by immunoblot analysis (Fig. 2d, e and Extended Data Fig. 4e, f). Furthermore, LAMP1, a lysosomal membrane marker, was redistributed from the perinuclear region to the cell periphery and dendritic processes, indicating enhanced membrane association (Fig. 2f, g and Extended Data Fig. 4g, h). To directly visualize secretory lysosomes in live cells, we employed total internal reflection fluorescence (TIRF) microscopy ^18–20^. NTR astrocytes labeled with FITC-Dextran and lysosome-RFP exhibited frequent exocytic events near the plasma membrane following treatment with α-syn PFF-activated MCM (Fig. 2h, i and Extended Data Fig. 4i). These events were characterized by transient brightening and dimming at the cell surface, consistent with partial release of lysosomal contents rather than full organelle extrusion.

The machinery driving the secretion of lysosomal contents involves coordinated transport and membrane fusion steps mediated by VAMP7, SNAP23, TRPML1, and synaptotagmin-7 (Syt-VII) ^21,22^. To determine the functional relevance of this machinery in NTR astrocytes, we used lentiviral CRISPR/Cas9-mediated knockdown of these genes in primary cultured astrocytes (Extended Data Fig. 5a-e). Loss of each of these components significantly reduced lysosomal contents release and impaired peripheral localization of LAMP1 (Fig. 2j, k and Extended Data Fig. 5f, g). Together, these findings demonstrate that NTR astrocytes exhibit active lysosomal exocytosis, releasing acid hydrolases and catabolic enzymes into the extracellular milieu.

### NTR astrocytes acidify the extracellular microenvironment

Lysosomal exocytosis releases protons (H⁺) into the extracellular space, thereby acidifying the surrounding microenvironment ^23–25^. To investigate whether NTR astrocytes contribute to extracellular acidification in the brain, the extracellular pH of α-syn PFF-induced NTR astrocytes or quiescent astrocytes was measured using a pH meter and an extracellular pH sensor (Fig. 3a). Both assays confirmed that the supernatants from α-syn PFF-induced NTR astrocytes was more acidic than those from quiescent astrocytes (Fig. 3b, c). NTR astrocytes directly induced by IL-1α, TNFα and C1q also showed similar extracellular pH reduction (Fig. 3d). Inhibition of lysosomal exocytosis-related genes markedly suppressed extracellular acidification of astrocytes induced by α-syn PFF-MCM, suggesting a direct link between lysosomal exocytosis and extracellular acidification (Fig. 3e).

**Fig. 3.**
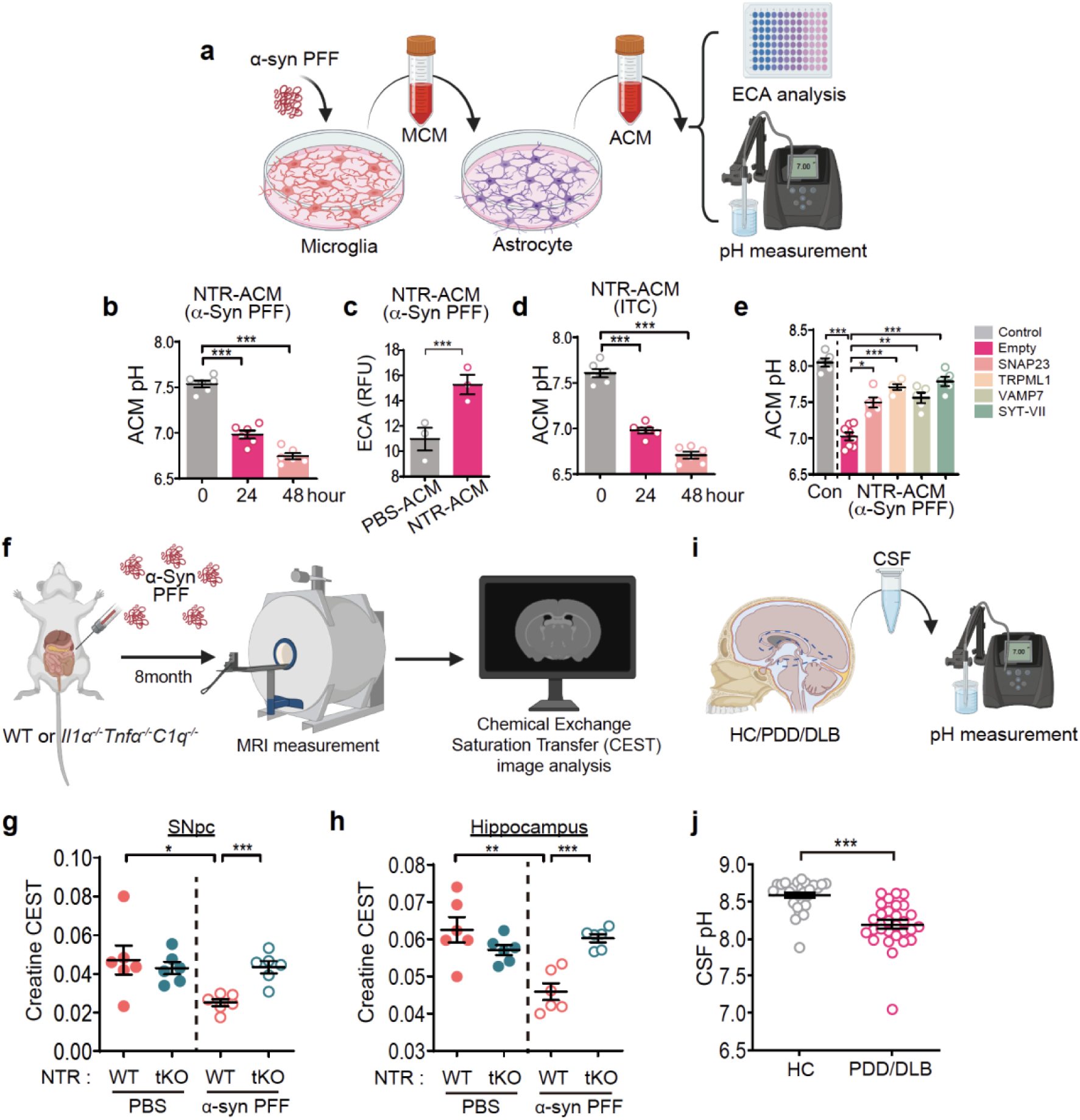
Acidification on extracellular space induced by NTR astrocyte lysosomal exocytosis. **(a)** Schematic of the conditioned media treatment. Microglia conditioned media (MCM) from PBS or α-syn PFF treated primary microglia was collected and added to cultured primary astrocytes and the acidity level was assessed in the astrocyte conditioned media (ACM). The figure was created with BioRender. **(b)** NTR-ACM pH after treatment with MCM derived from treatment of microglia with PBS (PBS-MCM) or α-syn PFF (PFF-MCM) at 0, 24, 48 h. . Data are the means ± s.e.m. Unpaired two-tailed student *t* test (n=6). **(c)** Extracellular acidification (ECA) determined as an extracellular acidification assay kit used to quantify changes in proton release and pH in the extracellular environment. extracellular pH sensor kit at 24 hours from NTR-ACM after treatment with PBS-MCM or PFF-MCM, two-tailed Student’s t test (n=3). Data are the mean ± s.e.m. p < 0.05; **p < 0.01; ***p < 0.001, two-tailed Student’s t test. **(d)** NTR-ACM pH after treatment with PBS-MCM or Il1α, TNFα, C1q – MCM at 0, 24, 48h. Data are the means ± s.e.m. Unpaired two-tailed student *t* test (n=6). **(e)** NTR-ACM pH 24 h after treatment with PBS-MCM (Cont) or PFF-MCM from astrocytes transfected with lentiCRISPR-v2 Mock and SNAP23, TRPML1, VAMP7, SYT-VII prior to treatment with PBS-MCM or PFF-MCM. Data are the means ± s.e.m. Two-way ANOVA followed by Bonferroni’s post hoc test (n=5-8). **(f)** Schematic of MRI scan images determined 8 months after PBS or α-syn PFF gut injection in WT (n=6) and tKO mice (n=6). **(g, h)** MRI creatine CEST values in (g) substantia nigra and (h) hippocampus. Data are the mean ± s.e.m. Two-way ANOVA followed by Bonferroni’s post hoc test (n=6). **(i, j)** LBD (DLB and PDD) patients CSF pH (n=29 for HC, n=29 for LBD patients). All data were analyzed by two-way ANOVA followed by post hoc Bonferroni test for multiple group comparison. Error bars represent the mean ± s.e.m. *p < 0.05; **p < 0.01; ***p < 0.001.

To determine whether acidification occurs in vivo, creatine chemical exchange saturation transfer (CrCEST) magnetic resonance imaging (MRI) which can be used to monitor extracellular pH of brains ^26^ was performed 8 months after α-syn PFF injection into the gut of WT and tKO mice (Fig. 3f). CrCEST signals were reduced in the SNc and hippocampus of WT mice post α-syn PFF injection while there was no change in these same brain regions in tKO mice post α-syn PFF injection (Fig. 3g, h). In addition, CSFs from Lewy body dementia (PDD and DLB) patients were significantly more acidic than those from age-matched healthy controls (Fig. 3i, j and Table S3). These results suggest that brain pH is reduced in the setting of pathologic α-syn-induced neurodegeneration in NTR astrocyte-dependent manner.

### NTR astrocyte-induced extracellular acidification drives neurotoxicity via Acid-sensing ion channel 1 (ASIC1)

Extracellular acidosis is a driver of neuronal dysfunction and degeneration, in part through the activation of acid-sensing ion channels (ASICs) such as ASIC1a ^27–31^. ASIC1a is abundantly expressed in the central nervous system and upregulated under pathological conditions where it facilitates calcium influx in response to lowered pH and is implicated in promoting cell death and inflammatory signaling during neurodegenerative disease states ^32,33^. To determine the role of acidic microenvironment in pathologic α-syn-induced neurodegeneration, ASIC1 expression was monitored in MAP2^+^ neurons and found to be significantly upregulated in postmortem midbrain of DLB and PDD brains (Fig. 4a-c) and in the midbrain of the gut-to-brain α-synucleinopathy mouse model (Fig. 4d, e) as determined by immunohistochemistry. ASIC1 expression significantly increases in primary cortical neurons treated with α-syn PFF-induced NTR-ACM compared to control-ACM (Fig. 4f-h). ASIC1 expression also increased in culture media at pH of 6.8 that is observed in α-syn PFF-induced NTR-ACM (see Fig. 3b) (Fig. 4g, i). No change in the expression of ASIC2, 3 and 4 was observed under similar conditions (Extended Data Fig. 6a-n).

**Fig. 4.**
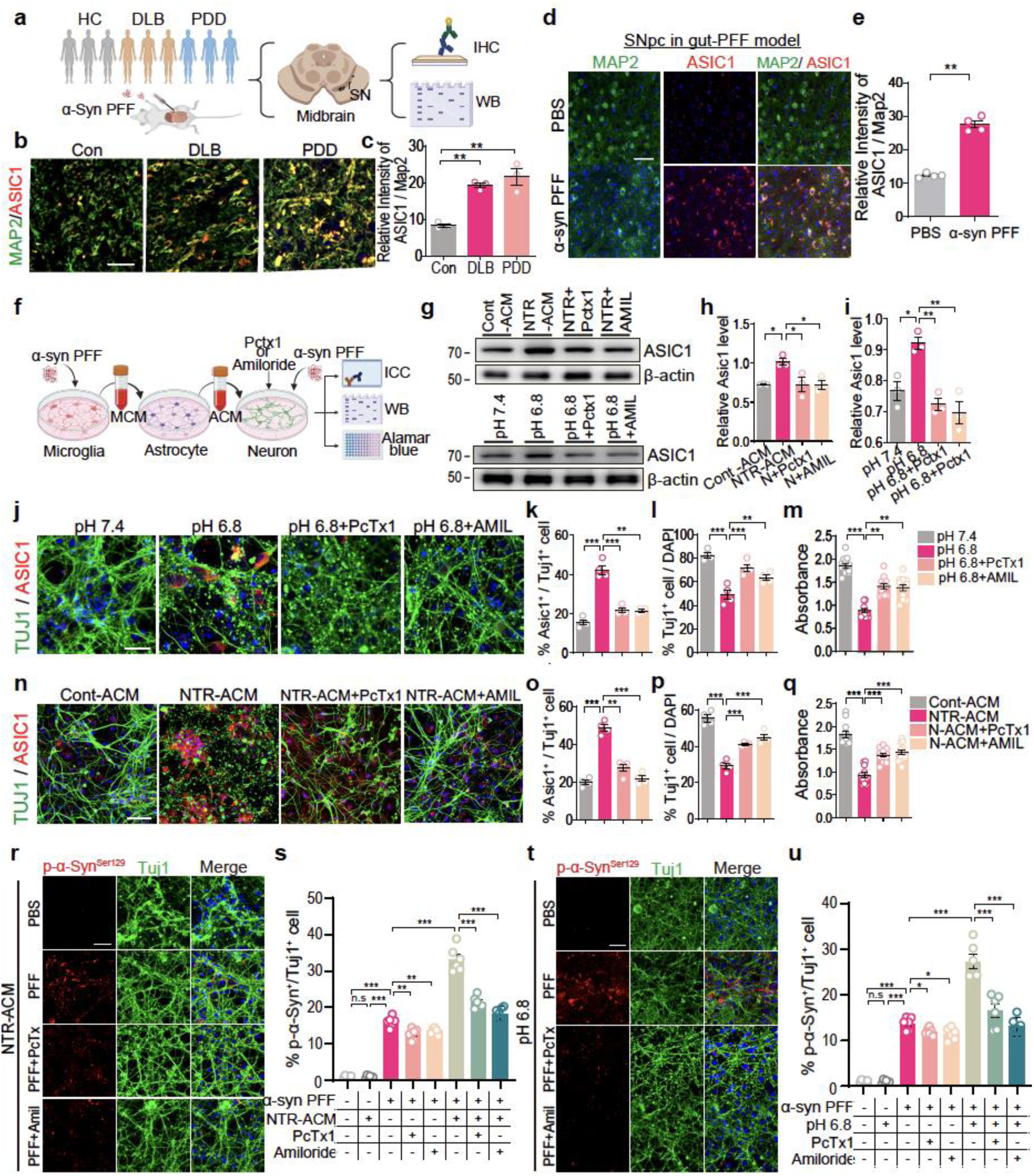
Asic1 level is increased in the brain of pathogenic environments and primary Neurons treated with NTR-ACM and acidic medium. **(a)** Schematic diagram of the biochemical analyses through gastrointestinal injection of α - syn PFF and human DLB and PDD postmortem samples. **(b)** Representative images of double immunostaining for Map2 (green) and ASIC1 (red) in the midbrain of DLB, PDD and control postmortem patient samples. Scale bars, 20 μm. **(c)** Quantification of relative intensity of ASIC1+ cells within MAP2+ cells (n=3). **(d)** Representative images of double immunostaining for MAP2 (green) and ASIC1 (red) in the midbrain 8 months after α-syn PFF gut injection in WT mice. Scale bars, 50 μm. **(e)** Quantification of the relative intensity of ASIC1+ cells within MAP2+ cells (n=4). **(f)** Schematic of the CM treatment experiments. Media conditioned in PBS or α-syn PFF treated primary microglia was collected and added to cultured primary astrocytes and analyze acidity level in the Astrocyte CM. **(g)** Representative Immunoblots of ASIC1 and β -actin of primary mouse cortical neurons in the top panel treated with PBS-MCM induced NTR-ACM (Cont), α-syn PFF-MCM induced NTR-ACM (NTR-ACM) and α-syn PFF-MCM induced NTR-ACM treated with ASIC1 inhibitor, PcTx1 (Psalmotoxin-1) treatment: 30 nM, 24h (N + PcTx1) and the bottom panel primary mouse cortical neurons in pH 7.4, pH 6.8 culture media and pH 6.8 + PcTx1 (30 nM) 24h culture media. PcTx1, a peptide toxin that selectively inhibits ASIC1a channels and applied to neuronal cultures at a concentration of 30 nM. **(h, i)** Quantification of ASIC1 normalized to β-actin from panel g. Data are the means ± s.e.m. Two-way ANOVA followed by Bonferroni’s post hoc test (n=4). **(j)** Representative immunofluorescence images of the neuronal marker (TUJ1, green) and ASIC1(red) in primary cortical neurons treated with Cont-ACM, NTR-ACM, NTR-ACM + PcTx1 and NTR-ACM + AMIL(Amiloride, 20 μM) for 24h (n=4). **(k)** Percentage of ASIC1^+^ cells within TUJ1^+^ cells (n=4). **(l)** Percentage of TUJ1^+^ cell with DAPI^+^ cells (n=4). **(m)** Cell viability was measured using Alamar blue absorbance. Data are the means ± s.e.m. Two-way ANOVA followed by Bonferroni’s post hoc test (n=10-12). **(n)** Representative immunofluorescence images of the neuronal marker (Tuj1, green) and ASIC1(red) in primary cortical neurons in pH 7.4 and pH 6.8 culture media and pH 6.8 culture media with PcTx1 or AMIL. **(o)** Percentage of ASIC1^+^ cells within TUJ1^+^ cells (n=4). **(p)** Percentage of TUJ1^+^ cell with DAPI^+^ cells (n=4). **(q)** Cell viability was measured using Alamar blue absorbance. Data are the means ± s.e.m. Two-way ANOVA followed by Bonferroni’s post hoc test (n=10-12). **(r-u)** Representative double-immunostaining for pSer129-α-syn (red) and TUJ1 (green) in primary cortical neurons induced by α-syn PFF with (r) NTR-ACM or (tr) acidic medium (pH 6.8) treated with vehicle or PcTx1 or Amiloride, (n=6, biologically independent primary cortical neurons). Scale bar, 20 μm. Percentage of TUJ1 positive neurons with pSer129-α-syn (p-α-syn) of cortical neurons induced by α-syn PFF with (s) NTR-ACM or (u) acidic medium (pH 6.8) treated with vehicle or PcTx1 or Amiloride, Bars represent the mean ± S.E.M. Two-way ANOVA followed by Bonferroni’s post hoc test (n=6, biologically independent primary cortical neurons). p < 0.05; **p < 0.01; ***p < 0.001.

To determine whether ASIC1 contributes to reactive astrocyte-induced neurotoxicity in response to acidic extracellular milieu, mouse primary cortical neurons were cultured in acidic medium or treated with NTR-ACM. Neurons exposed to acidic environment by either acidic medium or NTR-ACM exhibited significant cell death, as measured by TUJ1 immunostaining and the Alamar Blue assay (Fig. 4j-q). Treatment with ASIC1 inhibitors, PcTx1 ^32,34,35^ or amiloride ^32,36,37^, suppressed the increased expression of ASIC1 (Fig. 4j, k, n, o) and reactive astrocyte-induced neurotoxicity (Fig. 4l, m, p, q). Suppression of lysosomal exocytosis of NTR astrocyte failed to induce ASIC1 expression in neurons and NTR-astrocyte-induced neurotoxicity (Extended Data Fig. 7a-f). In addition, pharmacological inhibition of ASIC1 suppressed reactive astrocyte-induced synaptic degeneration in both mouse primary cortical and dopaminergic neuron cultures (Extended Data Fig. 8a-f).

Lentiviral CRISPR/Cas9-mediated knockdown of ASIC1, but not ASIC2-4, showed neuroprotection against NTR-ACM treatment, evidenced by increased neuronal survival and preserved neurite length (Extended Data Fig. 8g-j). Given that ASIC1 activation elevates intracellular calcium levels ^35,37^ and that calcium dysregulation is closely associated with neuronal damage ^38,39^, we found that NTR-ACM induced significant Ca^2+^ influx in primary cultured neurons, which was prevented by treatment of ASIC1 inhibitors (Extended Data Fig. 8k).

To ascertain whether extracellular acidification could influence α-synuclein pathology in neurons, primary cortical neurons were pretreated with PBS or a non-lethal dose of α-syn PFFs for 7 days and then exposed to NTR-ACM or acidic media, with or without ASIC1 inhibitors. Pathologic α-syn accumulation measured by pSer129-α-syn immunostaining and immunoblots of soluble and insoluble α-syn species were significantly elevated following NTR-ACM or acidic medium exposure that was attenuated by ASIC1 inhibition (Fig. 4r-u and Extended Data Fig. 9a-e). Taken together, these data implicate ASIC1 as a substantial mediator of neuronal vulnerability to NTR astrocyte-derived acidosis and that extracellular acidosis not only promotes neuronal death but also promotes pathologic α-synuclein accumulation in neurons.

### Genetic deletion or pharmacological inhibition of ASIC1a ameliorates behavioral deficits and neuropathological features in vivo

To check the therapeutic potential of ASIC1 inhibition, we evaluated whether genetic deletion or pharmacological blockade of ASIC1 could alleviate α-synucleinopathy-associated pathology in the gut-to-brain α-synucleinopathy mouse model where α-syn PFFs were injected in the gut of *Asic1a^f^*^lox/flox^;CaMKII-Cre conditional knockout mice (ASIC1a cKO), which lack neuronal ASIC1 expression, and in WT mice with oral administration of the ASIC1 inhibitor, amiloride (Extended Data Fig. 10a-c).

Behavioral tests revealed that gut-PFF-injected littermate control mice (*Asic1a*^flox/flox^; CaMKII-Cre^-/-^) (Fig. 5a) developed motor impairments 8 months after the gut-PFF injection as assessed by pole test, grip strength, open field tests (Fig. 5b, c and Extended Data Fig. 10d-i) and cognitive deficits as assessed by Morris water maze and Y-maze tests (Fig. 5d, e and Extended Data Fig. 10j-n). On the other hand, ASIC1a cKO mice injected with α-syn PFFs exhibited significantly improved performance across all behavioral measures compared to their littermate controls (Fig. 5b-e and Extended Data Fig. 10d-n). Neurodegeneration of DA neurons was assessed via TH immunostaining followed by stereological analysis, TH immunoblotting and measurement of striatal DA levels. There was significant DA neuronal loss in littermate control mice after gut-PFF injection. In contrast, ASIC1a deletion preserved TH⁺ and Nissl⁺ neuron numbers and striatal DA levels (Fig. 5f-h and Extended Data Fig. 10o-s). Synaptic integrity was evaluated by measuring the levels of synaptophysin and PSD95 in the hippocampus. WT mice injected with α-syn PFFs showed a significant reduction in these markers while ASIC1a deletion prevented the loss of these synaptic markers (Fig. 5i-k).

**Fig. 5.**
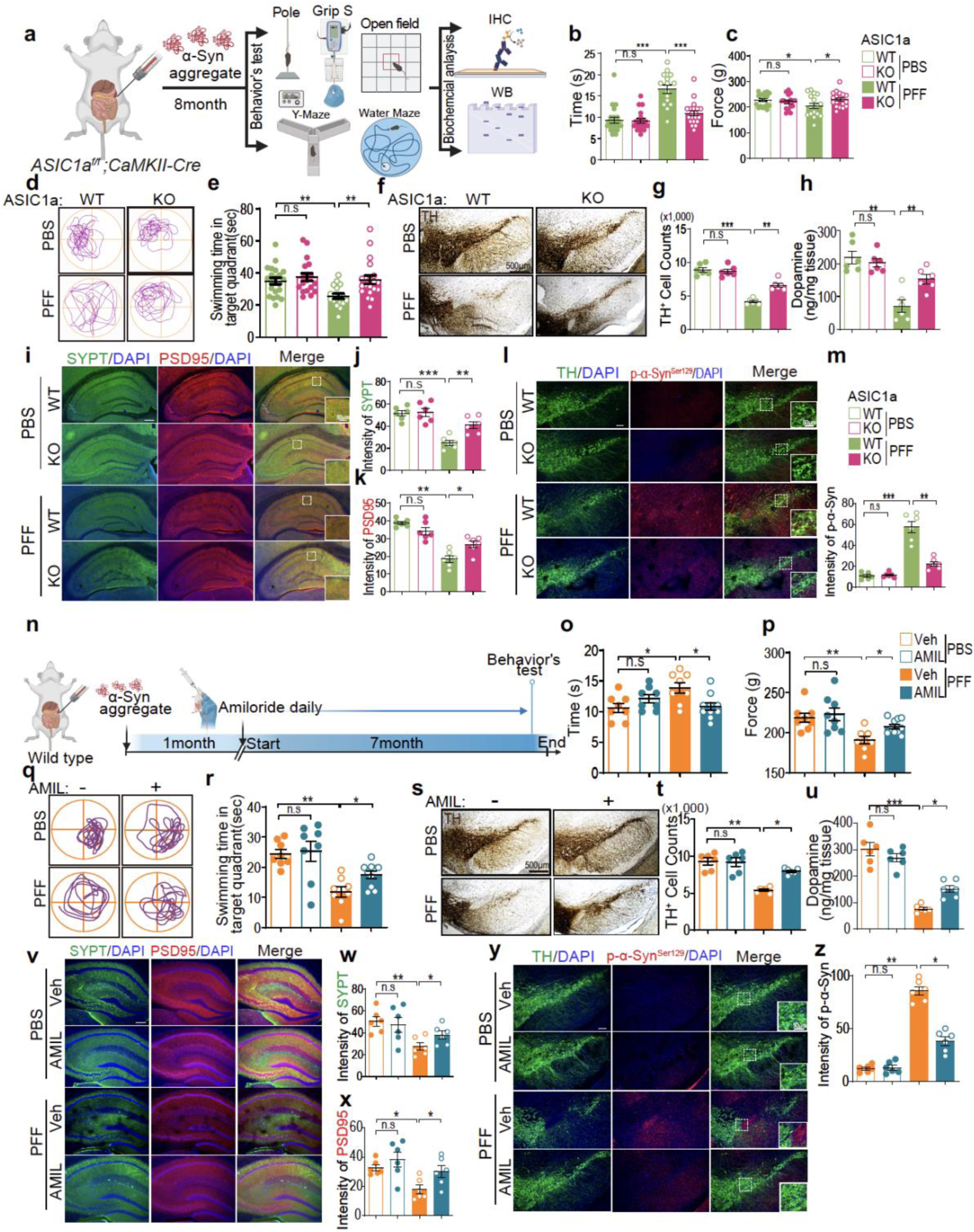
DLB and PDD-like pathology in the gut-to-brain α -syn PFF model is rescued by genetic deletion of neuronal ASIC1a or inhibition of ASIC1. **(a)** Schematic diagram of Motor function and cognitive behavioral assessments 8 months after PBS and α-syn PFF gastrointestinal injection in CaMKII-Cre^-^; ASIC1a^flox/flox^ mice (Control) (n = 17-21), CaMKII-Cre^+^;ASIC1a^flox/flox^(Knockout) (n = 18-20). **(b, c)** Results of mice on the (b) turn and descend time of pole test, (c) fore and hindlimb force of grip strength test. **(d, e)** (d) Representative swimming paths of mice from each group in the Morris water maze test (MWMT) on the probe trial day 5. (e) Result of mice on the probe trial session in the Morris water maze test. **(f-h)** (f) Representative photomicrographs from coronal mesencephalon sections containing TH-positive neurons in the SNc region 9 months after PBS and α-syn PFF gastrointestinal injection (n = 18-20). Scale bar, 500 μm. e,f, Unbiased stereological counts of (g) TH-positive in the SNc region and (h) DA level of striatum(ng/mg of tissue) in CaMKII-Cre^-^; ASIC1a^flox/flox^ mice(Control) (n=17-21), CaMKII-Cre^+^;ASIC1a^flox/flox^ (KO). **(i-k)** (i) Representative immunofluorescence images of (j) Pre-synaptic marker (SYPT, green) and (k) Post-synaptic marker (PSD95, red) in the hippocampal region of CaMKII-Cre^-^; ASIC1a^flox/flox^ mice (Control) (n=6), CaMKII-Cre^+^;ASIC1a^flox/flox^ (KO) (n = 6).Scale bar, 200 μm. **(l-m)** (l) Representative double immunostaining for pSer129-α-syn (red) and TH (green) in the SNc of CaMKII-Cre^-^; ASIC1a^flox/flox^ mice (Control) (n=17-21), CaMKII-Cre^+^;ASIC1a^flox/flox^ (KO) (n=6). White dashed box in the left panel of outlines area of high magnification in the panels shown to the right. n = 6 biologically independent animals. Scale bar, 100 μm. (m) Intensity of pSer129-α-syn. Error bars represent the mean ± SEM. All biochemical analyzes and behavior tests were analyzed by two-way ANOVA followed by post hoc Bonferroni test for multiple group comparison. *P < 0.05, **P < 0.01, ***P < 0.001,. NS, not significant. **(n)** Schematic diagram of experimental design in gut-PFF model with oral administration of amiloride. Motor function and cognitive behavioral assessments 7 months after gut injection of α-syn PFF between mice treated with vehicle (PBS) and mice receiving daily oral injections of 5-10 mg kg^-1^ Amiloride (n=8-10). **(o-r)** Results of mice on the (o) turn and descend time of pole test, (p) fore and hindlimb force of grip strength test. (q) Representative swimming paths of mice from each group in the Morris water maze test (MWMT) on the probe trial day 5. (r) Result of mice on the probe trial session in the Morris water maze test. **(s-u)** Representative photomicrographs from coronal mesencephalon sections containing TH-positive neurons in the SNc region of Vehicle and Amiloride treated mice at 7 months after gut injection of α-syn PFF or PBS (n = 6). biologically independent animals. Scale bar, 500 μm. e,f, Unbiased stereological counts of (t) TH-positive in the SNc region, (u) DA level of striatum(ng/mg of tissue). Data are mean ± s.e.m.; n = 6, biologically independent animals. **(v-x)** (v) Representative immunofluorescence images of Pre-synaptic marker (Synaptophysin, green) and Post-synaptic marker (PSD95, red) in the hippocampal region of PBS and Amiloride treated mice at 6 months after gut injection of α-syn PFF or PBS. n = 6 biologically independent animals. Scale bar, 200 μm. e,f, intensity of (w) Pre, Synaptophysin and (x) Post, PSD95-positive synaptic cells in the hippocampal region of Vehicle and Amiloride treated mice at 6 months after gut injection of α-syn PFF or PBS. Data are mean ± s.e.m.; n = 6, biologically independent animals. **(y, z)** Representative double immunostaining for p-α -syn^Ser129^ (red) and TH (green) in the SNc. White dashed box in far left panel of outlines area of high magnification in the panels shown to the right. (n= 6). Scale bar, 100 μm. (z) Intensity of pSer129-α-syn in SNc of Vehicle and Amiloride treated mice at 6 months after gut injection of α-syn PFF or PBS. Data are mean ± s.e.m. (n=6). *P < 0.05, **P < 0.01, ***P < 0.001.

Immunostaining for pSer129-α-syn in the SNc, hippocampus, cortex and striatum of gut-PFF-injected littermate control mice and immunoblots of Triton X-soluble and -insoluble fractions in SNc region revealed elevated pathologic α-syn in the gut-PFF–injected littermate control mice while ASIC1a cKO significantly reduced pathologic α-syn accumulation (Fig. 5l, m and Extended Data Fig. 10t-x).

Oral administration of amiloride (5 mg/kg/day) was initiated one month after gut-PFF injection corresponding to the early phase of formation of NTR astrocytes (see Extended Data Fig. 1j-q) and continued daily for 7 months (Fig. 5n). To ensure brain bioavailability of amiloride, high-performance liquid chromatography (HPLC) in WT mice confirmed its penetration across the blood–brain barrier (Extended Data Fig. 11a, b). Behavioral, histological, and biochemical analyses were conducted 8 months post-injection of α-syn PFFs (Fig. 5n). Amiloride-treated mice exhibited significantly improved performance of motor (Fig. 5o, p and Extended Data Fig. 11c-h) and cognitive behavior (Fig. 5q, r and Extended Data Fig. 11i-l) compared to their respective controls injected with α-syn PFFs. In addition, WT mice treated with amiloride exhibited significantly less loss of DA neurons after the gut-PFF injection compared with mice treated with vehicle as assessed by stereological counting of TH immunoreactivity in the SNc (Fig. 5s, t). α-Syn-PFF-induced loss of TH immunoreactivity in the striatum and reduction in striatal DA levels were normalized by amiloride treatment (Fig. 5u and Extended Data Fig. 11m-o). In addition, α-syn-PFF-induced hippocampal synaptic loss was rescued in amiloride-treated mouse brain (Fig. 5v-x).

Pathological α-syn accumulation exhibited in WT mice was significantly attenuated in mice treated with amiloride (Fig. 5y, z and Extended Data Fig. 11p-v). Taken together, these results demonstrate that ASIC1 is an important mediator of pathologic α-syn induced neuronal degeneration and accompanying behavioral deficits.

## Discussion

This study identifies NTR astrocytes as pivotal contributors to neurodegeneration in α-synucleinopathies by promoting an acidic extracellular environment, which disrupts neuronal health and survival. While astrocyte reactivity plays a role in numerous neurodegenerative diseases ^40,41^, our findings refine the current paradigm by linking astrocyte reactivity—mediated by microglial cytokine signaling—to a cascade that decreases extracellular pH and compromises neuronal survival. This mechanism involves NTR astrocytes engaging in lysosomal exocytosis, actively releasing acidic and catabolic enzymes into the extracellular space. Proteomic analyses consistently revealed a lysosomal protein signature among NTR astrocytes induced by various neurotoxic stimuli, and live imaging verified enhanced lysosome secretion in these cells. Disruption of exocytosis machinery via knockdown of VAMP7, TRPML1, SNAP23, or SYT7 markedly reduced both lysosomal release and acidification, underscoring this pathway’s essential role in astrocyte-driven toxicity.

Measurable acidification was confirmed in the cerebrospinal fluid of DLB and PDD patients as well as in α-synucleinopathy mouse models, aligning with previous studies in Alzheimer’s disease models that reported reduced brain and CSF pH ^42^. Given that NTR astrocytes are also observed in other proteinopathies ^43,44^, it is plausible that astrocyte-driven acidification could be a general mechanism contributing to neuronal vulnerability in diverse neurodegenerative conditions. Further investigation into pH dynamics in human tissue and fluid specimens is important for validation.

On the neuronal side, the acidified environment produced by NTRs activates voltage-independent, proton-sensitive ASIC1 ion channels, which are highly responsive to pH values observed in disease states ^45^. Only ASIC1, not related channels, mediates the neurotoxic response to acidification: its activation enhances calcium influx and drives both synaptic degeneration and pathological α-synuclein accumulation, ultimately leading to neuronal death. Genetic or pharmacological blockade of ASIC1—using either neuron-specific knockout strategies or the diuretic amiloride^46^—was highly effective in protecting against α-synuclein-induced neurodegeneration, preserving both motor and cognitive functions, as well as neuronal and synaptic markers. Our findings overturn conventional models by showing that these astrocytes actively acidify the brain’s extracellular milieu via lysosomal exocytosis, thereby triggering neuronal death through proton-gated ASIC1 channels. This establishes an actionable mechanistic link between glial inflammation, extracellular acidification, and synaptic and neuronal loss. By identifying astrocyte-driven pH dynamics as a crucial mediator of disease progression, our work opens entirely new avenues for therapeutic intervention targeting astrocyte biology in dementia.

## Materials and Method

### Animals

All experimental procedures were in accordance with the guidelines of the Laboratory Animal Manual of the National Institute of Health Guide to the Care and Use of Animals, which were approved by the Johns Hopkins Medical Institute Animal Care and Use Committee. All housing, breeding, and procedures were performed according to the NIH Guide for the Care and Use of Experimental Animals and approved by Johns Hopkins University Animal Care and Use Committee.

#### Mouse strain for Intestinal intramuscular α-syn Gut-PFF injection

C57BL/6 mice were obtained from the Jackson Laboratories (Bar Harbor, ME, USA). The mice do not develop any autoimmune or inflammatory phenotype. Amiloride was subcutaneously administered (5-10mg/kg) daily one month after α-syn PFF stereotaxic injection until ten months.

#### Neurotoxic reactive astrocyte knockout mice

Neurotoxic reactive astrocyte triple knockout (*Il1a*^−/−^ *Tnfa*^−/−^*C1q*^−/−^) animals were obtained from the Dr. Shane A Liddelow (Neuroscience Institute, NYU School of Medicine, New York, NY, USA).

#### Acid-sensing ion channel 1a knockout mice

ASIC1a^loxP/loxP^ strain animals were obtained from the Dr. John A. Wemmie (University of Iowa, Iowa City, Iowa, USA) and intercrossed with CaMKII-Cre mice to produce neuronal ASIC1a knockout mice.

### α-Synuclein purification and α-syn PFF preparation

Recombinant mouse α-synuclein protein were prepared from IPTG independent inducible pRK172 vector ^47^. Bacterial pellets were resuspended in high-salt buffer [750 mM NaCl, 10 mM Tris (pH 7.6) and 1 mM EDTA] containing a mixture of protease inhibitors (Roche), boiled for 15 min, and centrifuged at 6,000 x g for 20 min. The supernatants were applied onto a Superdex 200column (GE Healthcare) and Hi-Trap Q HP anion-exchange column (GE Healthcare) and eluted at ∼300 mM NaCl. After purification, bacterial endotoxins were eliminated by Toxineraser endotoxin removal kit (GeneScript). α-syn PFF were prepared as previously described ^48^ by agitating (1,000 rpm at 37ᵒ C) for 7 days and sonicated for 30 s (0.5 sec pulse on/off) at 10% amplitude (Branson Digital Sonifier).

### Intestinal intramuscular α-syn PFF injection

α-syn PFF injection to intestinal intramuscular was conducted as previously described ^14^. Mice at 3 months of age were anesthetized using avertin and kept at a constant body temperature using a conventional heat pad. For each animal, the injection was conducted using a 10 mL Hamilton syringe into the wall of the pyloric stomach at 2 sites and intestine wall of the duodenum at 2 sites, 0.5 cm apart from the pyloric stomach injection site.

Injections were made near the myenteric plexus. The pyloric stomach was injected with 6.25 μg α-syn PFF in two different locations (2.5 mg/ml, 2.5 μl/location) for a total of 12. 5 μg α-syn PFF, and the upper duodenum was injected with 6.25 μg α-syn PFF in two different locations (2.5 mg/ml, total 2.5 μl/location) for a total of 12.5 μg α-syn PFF. In addition, control mice were injected with an equivalent volume of PBS at the same locations.

Following the injections, the animals were sutured and returned to normal housing conditions. At 2, 6, 8, and 10 months after α-syn PFF injection, the mice were sacrificed.

### Secretome analysis using Liquid Chromatography-Mass Spectrometry

#### Sample preparation

Sample preparation was conducted as previously described with minor modifications ^49,50^. One milliliter of culture medium containing secreted proteins derived from mouse astrocytes was used as the starting sample. Urea powder and TEAB solution were directly added to this sample to prepare a final volume of 1.8 mL, achieving final concentrations of 8 M urea and 50 mM TEAB. The samples were then reduced and alkylated with 10 mM tris(2-carboxyethyl)phosphine (TCEP, Sigma-Aldrich) and 40 mM 2-chloroacetamide (CAA, Sigma-Aldrich) at room temperature (RT) for 1 hour. They were concentrated to below 30 µL using Amicon Ultracel 30K filters (EMD Millipore) by centrifugation at 140,000 × g at 4 °C and subsequently concentrated again to 100 µL by adding 600 µL of lysis buffer. Secreted proteins were digested by adding 10 ng/µL of LysC (Lysyl endopeptidase; Fujifilm Wako Pure Chemical Industries Co., Ltd.) to each sample on the filter, followed by incubation at 37 °C for 3 hours. The samples were then transferred to 1.5 mL tubes. To dilute the urea concentration to 2 M, three volumes of 50 mM TEAB were added to the filter, and the resulting solution was also transferred to the same tube to recover any remaining sample. Further digestion was conducted by adding 10 ng/µL of trypsin (sequencing grade modified trypsin; Promega) and incubating at 37 °C overnight. Peptides were desalted using SCX StageTips by inserting a strong cation exchange disk (CDS Empore) into a 200 μL pipette tip. After peptide quantification using the Pierce™ Quantitative Peptide Colorimetric Assay Kit (Thermo Fisher Scientific), the samples were labeled with 6-plex TMT reagents (Thermo Fisher Scientific) according to the manufacturer’s instructions. Following labeling, 1/10 volume of 1 M Tris buffer was added to each sample to quench any remaining TMT reagents.

#### Peptide Fractionation Using SCX Stage Tip

Peptide fractionation was performed using SCX StageTips. TMT-labeled samples were adjusted to a final volume of 200 µL by adding 50 mM TEAB and then acidified by adding trifluoroacetic acid (TFA) to a final concentration of 1% before fractionation. Each SCX StageTip was activated with 200 µL of 100% acetonitrile (ACN) by centrifugation at 500 × g at RT. Samples were loaded onto the SCX StageTips by centrifugation at 100 × g at RT. The tips were washed with a total of 900 µL of 0.2% TFA by centrifugation at 1,000 × g at RT. Peptides were fractionated by sequential elution using six elution buffers. The first through fifth elution buffers consisted of 20% ACN and 0.5% formic acid (FA), each containing increasing concentrations of ammonium acetate: 50, 75, 125, 200, and 300 mM. The final fraction was eluted with a sixth elution buffer, Buffer X, consisting of 5% ammonium hydroxide and 80% ACN. At each elution step, 100 μL of elution buffer was applied to the SCX StageTip, and eluates were collected by centrifugation at 100 × g at RT. All fractions were dried using a vacuum concentrator and stored at –80°C until further analysis.

#### LC-MS/MS analysis

Peptide fractions were reconstituted in 49 μL of 0.5% FA, and 15 μL was injected in triplicate into the instrument for LC-MS/MS analysis. The Orbitrap Fusion Lumos Tribrid Mass Spectrometer (Thermo Fisher Scientific) coupled with an Ultimate 3000 RSLCnano nanoflow liquid chromatography system (Thermo Fisher Scientific) was used to analyze fractionated peptides. The peptides from each fraction were loaded onto a trap column (Acclaim PepMap 100, LC C18, 5 μm, 100 μm × 2 cm, nanoViper) at a flow rate of 8 μL/min, and then resolved at a flow rate of 0.3 μL/min by increasing the gradient of solvent B (0.1% FA in 95% ACN) to 28% on an analytical column (Easy-Spray PepMap RSLC C18, 2 μm, 75 μm × 50 cm) fitted onto an EASY-Spray ion source that was operated at a voltage of about 2.5 kV. The overall run time was 120 min. MS analysis was performed in data-dependent acquisition and “Top Speed” modes with 3 s per cycle. MS1 scan range for precursor ions was set to *m/z* 300 to 1,800. MS1 and MS2 mass resolution was set to 120,000 and 50,000 at an *m/z* of 200. MS2 scans were acquired by fragmenting precursor ions using the higher-energy collisional dissociation (HCD) mode, which was set to 35% of normalized collision energy. The automatic gain controls for MS1 and MS2 were set to one million and 0.05 million ions, respectively. The maximum ion injection times for MS1 and MS2 were set to 50 and 100 ms, respectively. The precursor isolation window was set to 1.6 with 0.4 of offset. Dynamic exclusion was set to 30 s, and singly charged ions were rejected. Internal calibration was carried out using the lock mass option (*m/z* 445.12002) from ambient air.

#### Database searches

Database searches and statistical analysis were conducted as described previously with minor modifications ^51^. Proteome Discoverer (version 2.4; Thermo Fisher Scientific) software was used for quantitation and identification. The top ten peaks in each window of 100 Da were selected for database search during MS2 preprocessing. The MS/MS data were then searched using SEQUEST HT algorithms against a Mus musculus UniProt database that includes both Swiss-Prot and TrEMBL (released in January 2019 with 55,435 entries) with common contaminant proteins (115 entries) ^52^. During MS2 preprocessing, the top ten peaks within each 100 m/z window were chosen for database search. The following parameters were applied for the database search: Trypsin was designated as the protease, allowing for a maximum of two missed cleavages. Fixed modifications included carbamidomethylation of cysteine (+57.02146 Da) and TMT tags (+229.16293 Da) on lysine and peptide N termini. Methionine oxidation (+15.99492 Da) was considered as a variable modification. A minimum peptide length of six amino acids was set. Precursor mass (MS1) and fragment mass (MS2) tolerances were established at 10 ppm and 20 ppm, respectively. False discovery rate (FDR) filtering was applied at 1% for both peptides and proteins using the percolator node and protein FDR validator node. For protein quantification, the integration mode employed the most confident centroid option, with a reporter ion tolerance of 20 ppm. The MS order was set to MS2, and HCD was selected as the activation type. Peptide quantification used both unique and razor peptides, while protein groups were considered for peptide uniqueness. The coisolation threshold was set at 50%. Reporter ion abundance calculations were based on signal-to-noise ratios, with missing intensity values replaced by the minimum value. An average reporter signal-to-noise threshold of 10 was applied. Corrections for isobaric tags and data normalization were disabled. Protein grouping followed a strict parsimony principle, grouping proteins sharing the same set or subset of identified peptides. Protein groups lacking unique peptides were filtered out. The final protein groups were generated by iterating through all spectra, selecting PSMs with the highest number of unambiguous and unique peptides, and summing the reporter ion abundances of PSMs for corresponding proteins

#### Statistical and bioinformatics analysis

The statistical analysis of the mass spectrometry data was performed with the Perseus software (version 1.6.0.7) ^53^. For normalization, reporter ion intensity values were divided by the median value of each protein, and the relative abundance values for each sample were log2-transformed. Subsequently, column-wise normalization was performed by subtracting the median value of each sample from its corresponding log2-transformed values. The P values between the comparison groups were calculated using the student’s two-sample t-test. Proteins with q-values < 0.05 were considered differentially expressed. Additional details about the number of samples (N) and replicates have been included in the figure legends. The q-values for the volcano plot were calculated by significance analysis of microarray (SAM) and a permutation-based FDR estimation with 0.1 of the S0 value ^54^. PCA and heatmap analyses were conducted using the MetaboAnalyst tool ^55^. Proteins exhibiting the significant protein abundance difference were subjected to KEGG pathway analysis ^56^ embedded in Enrichr ^57^ for gene set enrichment analysis.

### Behavioral tests

The behavioral deficits in gastrointestinal injection of α-syn PFF injected WT or NTR-KO mice, in gastrointestinal injection of α-syn PFF injected mice fed Vehicle or Amiloride, in gastrointestinal injection of α-syn PFF injected CAMKII-Cre^-^; ASIC1a ^flox/flox^ or CAMKII-Cre^+^; ASIC1a ^flox/flox^ were assessed by the pole test, grip strength test for motor deficits, and spontaneous alternation behavior Y-maze test, Morris water maze test for non-motor deficits 1 week prior to sacrifice as the different cohorts. All the experiments were performed by investigators who are blind to genotypes or treatment condition and randomly allocated to groups.

#### Pole test

The 9 mm diameter pole is a 2.5 ft metal rod wrapped with bandage gauze. Before the actual test, the mice were trained for two consecutive days and each training session consisted of three test trials. On the day of the test, mice were placed 3 inches from the top of the pole facing head-up. The time to turn and total time to reach the base of the pole were recorded. The maximum cutoff of time to stop the test and recording was 60 s.

#### Grip strength test

Neuromuscular strength was measured by maximum holding force developed by the mice using a grip-strength meter (Bioseb). Mice were placed onto a metal grid to grasp with either fore or both limbs that are recorded as ‘fore limb’ and ‘fore and hindlimb’, respectively. The tail was gently pulled, and the maximum holding force was recorded by the force transducer before the mice released their grasp on the grid. The peak holding strength was digitally recorded and displayed as force in grams (g).

#### Open field test

The open field consisted of a rectangular plastic box (40 cm X 40 cm X 40 cm) divided into 36 (6 X 6) identical sectors (6.6 cm X 6.6 cm). The field was subdivided into peripheral and central sectors, where the central sector included 4 central squares (2 X 2) and the peripheral sector was the remaining squares ^58^. The mouse was placed into the center of an open field and allowed to explore for 5 min under dim light. The apparatus was thoroughly cleaned with diluted 10% ethanol between each trial. A video tracking system (ANY-Maze software) was used to record the distance traveled as a measure of locomotor activity. The time spent in and entries into the center were measured as an anxiolytic indicator ^58^.

#### Y-maze test

A spontaneous alternation behavior Y-maze test was performed as described ^59^. The Y-maze is a horizontal maze with three equal angles between all arms, which were 40 cm long and 10 cm wide with 15 cm high walls. The maze floor and walls were constructed using opaque polyvinyl plastic. Mice were initially placed within one arm, and the sequence and number of arm entries were recorded manually for each mouse over an 8-min period. A spontaneous alternation was defined in which the mice entered all three arms, i.e., ABC, CAB, or BCA but not ABB, was recorded as an alternation to precision short-term memory. The alternation score (%) for each mouse was calculated as the ratio of the actual number of alternations to the possible number (defined as the total number of arm entries minus two) multiplied by 100 as shown by the following equation: % Alternative behavior = [(Number of alternations)/(Total arm entries – 2)] X 100. The number of arm entries per trial was used as an indicator of locomotor activity. The Y-maze arms were cleaned with diluted 10% ethanol between tests to eliminate odors and residues.

#### Morris water maze test (MWMT)

The MWMT was performed as described (Vorhees and Williams, 2006). The MWM is a white circular pool (150 cm in diameter and 50 cm in height) with four different inner cues on surface. The circular pool was filled with water and a nontoxic water-soluble white dye (20 ± 1℃) and the platform was submerged 1 cm below the surface of water so that it was invisible at water level. The pool was divided into four quadrants of equal area. A black platform (9 cm in diameter and 15 cm in height) was centered in one of the four quadrants of the pool. The location of each swimming mouse, from the start position to the platform, was digitized by a video tracking system (ANY-Maze, Stoelting Co., Wood Dale, IL, USA). The day before the experiment mice were subjected to swim training for 60 s in the absence of the platform. The mice were then given two trial sessions each day for four consecutive days, with an inter-trial interval of 15 min, and the escape latencies were recorded. This parameter was averaged for each session of trials and for each mouse. Once the mouse located the platform, it was permitted to remain on it for 10 s. If the mouse was unable to locate the platform within 60 s, it was placed on the platform for 10 s and then returned to its cage by the experimenter. On day 6, the probe trial test involved removing the platform from the pool and mice were allowed the cut-off time of 60 s.

### Dopamine ELISA

Extracellular dopamine concentration was measured with Dopamine ELISA (Cat. No. BA-E-5300R; Immusmol) following the manufacturer’s instructions. Briefly, mice were euthanized, and brains were rapidly extracted and placed on ice. The striatum was dissected bilaterally and homogenized in ice-cold phosphate-buffered saline (PBS) containing 0.1% Triton X-100 and protease inhibitors. Homogenates were centrifuged at 10,000 × g for 15 minutes at 4 °C to remove cellular debris, and the supernatants were collected for analysis. Prior to ELISA, samples underwent enzymatic derivatization to enhance dopamine detection sensitivity.

Dopamine ELISA was performed using a 750 µl volume of a sample. Each sample was a pool of 400 µl of striatum of brain lysates. Derivatized samples, standards, and controls were added to microtiter wells pre-coated with dopamine-specific antibodies and incubated to allow competitive binding. After washing, a secondary anti-rabbit IgG-peroxidase conjugate was applied, followed by tetramethylbenzidine (TMB) substrate for colorimetric detection. The reaction was terminated with stop solution, and absorbance was measured at 450 nm using a microplate reader. Dopamine concentrations were calculated from a standard curve using four-parameter logistic regression.

### Immunohistochemistry and immunofluorescence

Mice were perfused with PBS and 4 % PFA and brains were removed, followed by fixation in 4 % PFA overnight and transfer to 30 % sucrose for cryoprotection.

Immunohistochemistry (IHC) and immunofluorescence (IF) was performed on 40 μm thick serial brain sections. Primary antibodies and working dilutions are detailed in Table S4. For histological studies, free-floating sections were blocked with 10 % goat serum in PBS with 0.2 % Triton X-100 and incubated with TH or p-α-syn antibodies followed by incubation with biotin-conjugated anti-rabbit or mouse antibody, respectively. After three times of washing, ABC reagent (Vector laboratories, Burlingame, CA) was added and the sections were developed using SigmaFast DAB peroxidase substrate (Sigma-Aldrich). Sections were counterstained with Nissl (0.09 % thionin). For the quantification, both TH- and Nissl-positive DA neurons from the SNc region were counted by an investigator who was blind to genotypes or treatment condition with randomly allocated groups through optical fractionators, the unbiased method for cell counting, using a computer-assisted image analysis system consisting of an Axiophot photomicroscope (Carl Zeiss) equipped with a computer controlled motorized stage (Ludl Electronics, Hawthorne, NY), a Hitachi HV C20 camera, and Stereo Investigator software (MicroBright-Field, Williston, VT). The total number of TH-stained neurons and Nissl counts were analyzed as previously described ^60^. For immunofluorescent studies, double-labeled sections with TH and p-α-syn antibodies were incubated with a mixture of Alexa-fluor 488- and 594-conjugated secondary antibodies (Invitrogen). The fluorescent images were acquired by confocal scanning microscopy (LSM710, Carl Zeiss). All the images were processed by the Zen software (Carl Zeiss). The selected area in the signal intensity range of the threshold was measured using ImageJ software.

### Live imaging of lysosomal exocytosis

Cells were subjected to live imaging of cell membrane for injury-triggered lysosome assay as described ^61^. Cells were incubated overnight with growth media containing 2 mg/ml of FITC dextran. The cells were washed and incubated for 2 h with growth medium. Cells were then transferred to the microscope stage top incubator maintained at 37 °C. For live imaging, cells were imaged with TIRF at the penetration depth of 70–120 nm by imaging at for 30-60 min using Nikon iLas2 (Japanese multinational corporation, Tokyo, Japan) equipped with × 60/1.45 NA oil objective.

### Tissue- and cell-lysate preparation

Tissue lysates were prepared as described previously with some modifications ^62^. Non-ionic detergent-soluble and -insoluble fractions were made by homogenization of tissue in brain lysis buffer (10 mM Tris-HCl, pH 7.4, 150 mM NaCl, 5 mM EDTA, 0.5% Nonidet P-40, phosphatase inhibitor cocktail II and III (Sigma-Aldrich) and complete protease inhibitor mixture (Cell Signaling Technology, Danvers, MA, USA)). The homogenate was centrifuged at 22,000*g* for 20 min, and the resulting pellet and supernatant (soluble part) fractions were collected. The pellet was washed once in brain lysis buffer containing nonionic detergent (0.5% Nonidet P-40) and the resulting pellet (nonionic detergent insoluble) was homogenized in brain lysis buffer containing 1% SDS and 0.5% sodiumdeoxycholate. The homogenate was centrifuged and the resulting supernatant (nonionic detergent insoluble) was collected. Total cell lysates were prepared by homogenization of tissue in RIPA buffer (50 mM Tris, pH 8.0, 150 mM NaCl, 1% Nonidet P-40, 1% SDS, 0.5% sodium-deoxycholate, phosphatase inhibitor cocktail II and III (Sigma-Aldrich), and complete protease inhibitor mixture). The homogenate was centrifuged at 22,000*g* for 20 min, and the resulting supernatant (insoluble part) was collected.

### Immunoblot analysis

Mouse brain tissues and primary neurons were homogenized and prepared in lysis buffer. Primary antibodies and working dilutions are described in Supplementary Table 4.

Immunoblots of tissue and neuronal culture was performed as described previously ^62^. All raw blot and gel images are available in Supplementary Figs. 21, 22.

### Human postmortem brain tissues and cerebrospinal fluid

This research uses anonymous autopsy material; the substantia nigra from subjects with DLB/PDD and age-matched healthy controls from the Division of Neuropathology, Department of Pathology of Johns Hopkins University School of Medicine. The John Hopkins Medical Institutions Joint Committee on Clinical Investigations decided that these studies are exempt from Human Subjects Approval because of Federal Register 46.101 Exemption Number 4. Parkinson’s disease was confirmed by pathological and clinical criteria (Supplementary Table 1). NTR transcripts mRNA levels were monitored in human postmortem substantia nigra brain tissue from patients with Parkinson’s disease and controls (Supplementary Figure 3a) and immunohistochemistry. Cerebrospinal fluid (CSF) was collected by lumbar puncture at the Neurology clinics at Karolinska University Hospital, Sweden. Patients were included if they had a clinical diagnosis of DLB or PDD as determined by a neurologist. Control subjects were examined at the routine clinics for paraesthesia and headache with benign outcomes without any neurological condition or dementia. The study was approved by the Swedish Ethical Review Authority (Dnr the 2020-03684) and the Regional Ethical Review Board of Stockholm (2012/224-32/4).

### Primary neuron, microglia and astrocyte cell cultures, transfection and treatment

CD1 mice were obtained from Jackson Laboratories (Bar Harbor, ME, USA). Primary cortical neurons were prepared from embryonic day 15.5 pups and cultured in Neurobasal medium (Gibco) supplemented with B-27, 0.5 mM l-glutamine, penicillin and streptomycin (Invitrogen, Grand Island, NY, USA) on tissue-culture plates coated with poly-l-lysine. The neurons were maintained by changing the medium every 3–4 days. Primary microglial and astrocyte cultures were performed as described previously ^63^. Whole brains from mouse pups at postnatal day 1 (P1) were obtained. After removal of the meninges, the brains were washed in DMEM/F12 (Gibco) supplemented with 10% heat inactivated FBS, 50 U ml−1 penicillin, 50 μ g ml−1 streptomycin, 2 mM l-glutamine, 100 μ M non-essential amino acids and 2 Mm sodium pyruvate (DMEM/F12 complete medium) three times. The brains were transferred to 0.25% trypsin-EDTA followed by 10 min of gentle agitation. DMEM/F12 complete medium was used to stop the trypsinization. The brains were washed three times in this medium again. A single-cell suspension was obtained by trituration. Cell debris and aggregates were removed by passing the single-cell suspension through a 100-μm nylon mesh. The final single-cell suspension thus achieved was cultured in T75 flasks for 13 days, with a complete medium change on day 6. The mixed glial cell population was separated into astrocyte-rich and microglia-rich fractions using the EasySep Mouse CD11b Positive Selection Kit (StemCell). The magnetically separated fraction containing microglia and the pour-off fraction containing astrocytes were cultured separately. The conditioned medium from the primary microglia treated with α -syn PFF (α -syn PFF-MCM) with PBS treatment were collected and applied to primary astrocytes for 24 h. The conditioned medium from activated astrocytes by α-syn PFF-MCM, which we define as α-syn PFF-ACM, were collected with complete, Mini, EDTA-free Protease Inhibitor Cocktail (Sigma) and concentrated with Amicon Ultra-15 centrifugal filter unit (10 kDa cutoff) (Millipore) until approximately 50× concentrated. The total protein concentration was determined using Pierce BCA protein assay kit (Thermo Scientific), and 15 or 50 μg/ml of total protein was added to mouse primary neurons treated with α-syn PFF with acidic medium (pH 6.8), PcTx1, Amiloride treatment, respectively, for the neuronal cell death assay.

Primary neurons were infected with Lenti-control sgRNA or Lenti-ASIC1-4 gRNA at days in vitro (DIV) 4-5 by CRISPR/Cas9 strategy. The neuron growth medium was replaced with fresh medium. α-Syn PFF-ACM was added at DIV 7 and further incubated for indicated times followed by the cell death assay or biochemical experiments. Primary astrocytes were infected with Lenti-control sgRNA or Lenti-SNAP23, TRPML1, VAMP7, SYT-VII gRNA at DIV 5-6 and astrocyte growth medium was replaced with fresh medium every 2-3 days. α - syn PFF-MCM was added at DIV 9 and further incubated for indicated times followed by the measuring pH level or biochemical experiments. For co-culture experiments, mouse primary neurons were harvested and mixed with the mouse primary neurons astrocytes at a 2:1 ratio of neurons to astrocytes. The mixed cells were plated, and differentiation of mouse primary neurons was directly induced in serum-free N2 medium.

### Lentivirus construction and virus production

Mouse SNAP23, TRPML1, VAMP7, SYT-VII gRNA by CRISPR/Cas9 were subcloned into a lentiviral V2 vector ^64^. The lentivirus was produced by transient transfection of the recombinant cFugw vector into 293FT cells together with three packaging vectors: pLP1, pLP2, and pVSV-G (1.3:1.5:1:1.5). The viral supernatants were collected at 48 and 72 hours after transfection and concentrated by ultracentrifugation for 2 hours at 50,000 g.

### Morphometric assessments

To estimate morphological maturation, total Tuj1^+^ or Map2^+^ fiber length of 11 randomly selected Primary cultured cortical neurons were measured by Zeiss microscope equipped with automated computer assisted software (Axiovision 4.6, Carl Zeiss, Dublin, CA).

### The Alamar Blue and LDH assays

Cell death was assessed using the Alamar Blue assay (Invitrogen) according to the manufacturer’s protocol. LDH (Sigma) activity in the culture medium, representing relative cell viability and membrane integrity, was measured using the LDH assay kit spectrophotometrically, following the manufacturer’s instructions.

### pH measurements

Primary astrocytes were cultured in a modified DMEM medium devoid of buffering agents, including HEPES, sodium bicarbonate, and phenol red, to allow direct pH measurement without interference. The medium was prepared under sterile conditions and pre-warmed to 37 °C prior to use. All procedures were conducted in a CO₂-free environment to prevent pH fluctuations due to atmospheric CO₂. To assess extracellular pH changes, 1 mL of buffer-free medium was collected from each culture condition and transferred into sterile 1.5 mL microcentrifuge tubes. pH was measured using a calibrated microelectrode pH meter (PH8500-MS, Apera Instruments) equipped with a glass electrode suitable for small-volume samples. pH measurements were conducted at multiple time points, including baseline (pre-treatment)0, 24, 48 hour post-treatment of α -syn PFF-MCM or ITC cytokines (Il-1α,TNFα, C1q) and human CSF of control or DLP/PDD patients (Supplementary Table 3). All samples were measured in triplicate to ensure reproducibility. For drug-treated groups, astrocytes were exposed to the compound of interest following transfection, and medium was collected at designated intervals for pH analysis.

### Comparative qPCR

Total RNA from cultured cells was extracted with an RNA isolation kit (Qiagen, Valencia, CA, USA) following the instructions provided by the company. RNA concentration was measured spectrophotometrically using a NanoDrop 2000 (Biotek, Winooski, VT, USA). Subsequently, 1–2 μ g of total RNA was reverse transcribed to cDNA using the High Capacity cDNA Reverse Transcription System (Life Technologies, Grand Island, NY, USA). Comparative qPCR was performed in duplicate or triplicate for each sample using fast SYBRGreen Master Mix (Life Technologies) and ViiA 7 Real-Time PCR System (Applied Biosystems, Foster City, CA, USA). The expression levels of target genes were normalized to the expression of *Actb* and calculated based on the comparative cycle threshold *C*t method (2^−ΔΔ^*Ct*) .

### Measurement of Amiloride in brain

Amiloride were resuspended in a vehicle solution and were administered at 5-10 mg/kg by oral gavage. After 2 h, mice were perfused with PBS and whole brain was isolated, weighed and grounded with a pestle to a very fine powder with liquid nitrogen. Then, 2-folds of acetonitrile were added to brain tissue sample and the mixture was centrifuged at 14,000 rpm, 4℃ for 10 min. The supernatant was used for analysis. The concentration of compounds in brain tissue were measured using HPLC-MS on a C-18 reverse phase HPLC column.

Separations were achieved using a linear gradient of buffer B from 40% to 95% in A (A = 0.1% formic acid in H2O; B = 0.1% formic acid in CH3CN) at a flow rate of 1 ml/min.

Intraperitoneal (IP) injected brain samples at 0, 5, 10, 20 or 40 (mg/kg) concentration were used for the standard curve.

### MRI Experiments

All MRI experiments were performed on a horizontal bore 11.7 T Bruker Biospec system (Bruker, Ettlingen, Germany) equipped with actively shielded gradients with a maximum strength of 74 Gauss/cm. A 72 mm quadrature volume resonator was utilized for transmitting, and a 2 ×2 mouse phased array coil for detection. A double-tuned 31 P/ 1 H coil was employed for the collection of *in vivo* 31 P MRS. All coils were provided by Bruker. Anesthesia was induced using 2% isoflurane in medical air, followed by 1% to 1.5% isoflurane for maintenance during the MRI scan. The mouse head was positioned using a bite bar and two ear pins. During the MRI scan, mice were placed on a water-heated animal bed that was equipped with temperature and respiratory controls. The respiratory rate was monitored via a pressure sensor (SAII, Stony Brook, NY, USA) and maintained at 80-90 breaths per minute. The B 0 field over the mouse brain was adjusted using field mapping and second-order shimming. All CEST experiments were performed using continuous-wave CEST (cwCEST). This was described in detail in this paper ^26^.

### Calcium imaging

Measurement of intracellular calcium. Astrocytes were cultured on glass coverslips and incubated in DMEM with 10 μM of Fluo-4-AM (Life Technologies) for 20 min at 37 °C and 5% CO2. After washing with pre-warmed cell imaging media (CIM) imaged at four to eight frames/s. Change in cytosolic Ca2+ was measured as the ratio change in fluorescence over the background subtracted Fluo-4 fluorescence (ΔF/F0, F0 is Fluo-4 intensity before treatment and ΔF indicates the increase in fluorescence over F0).

## Cell counting and statistical analysis

Immunostained and DAPI-stained cells were counted in 9 to 20 random areas of each culture coverslip using an eyepiece grid at a magnification of ×200 or ×400. For every figure, data are expressed as the mean ± SEM and statistical tests are use as appropriate. Statistical comparisons were made using Student’s *t* test (unpaired), 2-tailed, or 1-way ANOVA followed by Bonferroni’s post hoc analysis using SPSS (Statistics 21; IBM Inc.). A *P* value of less than 0.05 was considered significant.

## Data availability

The mass spectrometry data from this study have been deposited to the ProteomeXchange Consortium (https://www.proteomexchange.org) via PRIDE partner repository with the dataset identifier PXD064107 and project name “Neurotoxic Reactive Astrocytes Drive Extracellular Acidification to Mediate α-Synuclein Neurodegeneration”. Reviewers can access the dataset only after logging into PRIDE using ‘’ as the ID and ‘pAQ63Kan5QRu’ as the password.

## Acknowledgements

The authors acknowledge the joint participation by the Adrienne Helis Malvin Medical Research Foundation through its direct engagement in the continuous active conduct of medical research in conjunction with The Johns Hopkins Hospital and the Johns Hopkins University School of Medicine and the Foundation’s Parkinson’s Disease Program H-2021. This work was supported by Parkinson’s Foundation, Freedom Together Foundation, the Robert J. and Claire Pasarow Foundation, the Knut and Alice Wallenberg Foundation and grants from the National Research Foundation of Korea (RS-2023-00278580, RS-2024-00407383) and Korea Dementia Research Center (RS-2024-00358266).

## Author Contributions

J.J.S., H.P. V.L.D., T.M.D and T.I.K. conceptualized the project. J.J.S., H.P. and T.I.K. designed the experiments. J.J.S. and H.P. performed experiments and analyzed data. J.J.S., H.P. Y.C., J.S., S.H.K. and S.H. performed immunoblotting and immunocytochemistry. J.J.S., H.P., A.P., J.W. and S.C.C performed behavioral experiments. J.J.S., H.P., A.P., D.B. and Y.S. performed histology and stereological cell counting. T.R., Y.J. and C.H.N. performed and analyzed mass spectrometry proteomics experiments. J.J.S and I.H. performed total internal reflection fluorescence (TIRF) microscopy experiments. J.W. generated and provided *ASIC1a^loxP/loxP^* mice. P.S. and J.T. collected and provided human CSFs and postmortem tissues. J.J.S. and J.X. performed creatine chemical exchange saturation transfer (CrCEST) MRI experiments. J.J.S., H.P., G.C. and T.I.K. prepared the figures. J.J.S., H.P., V.L.D, T.M.D. and T.I.K. wrote the manuscript, which all authors reviewed and edited. T.M.D, V.L.D, and T.I.K. acquired funding and supervised the project.

## Corresponding Author

Correspondence and requests for materials to Tae-In Kam.

## Competing interests

The authors declare no competing interests.

## Notes

### Competing Interest Statement

The authors have declared no competing interest.

### Summary of Updates

Typos updated; Supplemental files updated; Method section updated.

