## Supplemental Figures and Tables for "Reactive Astrocytes Drive Extracellular Acidification to Mediate α-Synuclein Neurodegeneration"

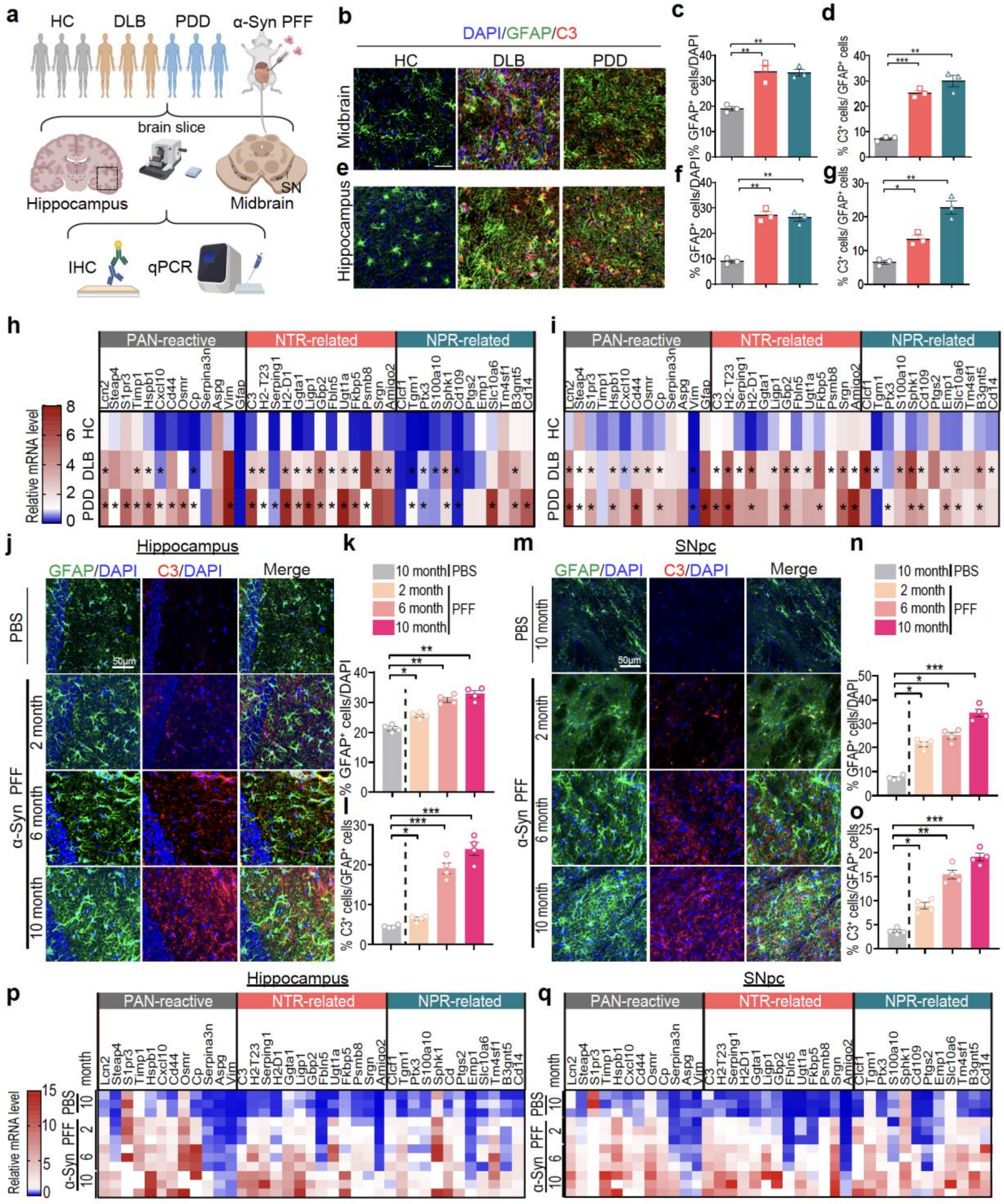


**Extended Data Figure 1. NTR astrocytes are increased in brain of human PDD and DLB patients and α-syn PFF gastrointestinal injection model.**

**(a)** Schematic diagram of the pathological and biochemical analyses through gastrointestinal injection of α-syn PFF and human DLB and PDD postmortem samples. The figure was created with BioRender.

**(b-g)** Representative images of double immunostaining for GFAP(green) and C3 (red) in (b) midbrain and (e) hippocampus in human DLB and PDD. Scale bars, 50 μm. Quantification of (c, f) percentage of $GFAP^+$ cells with DAPI and (d, g) $C3^+$ cells with $GFAP^+$ cells. Data are the means ± s.e.m. One-way ANOVA followed by Bonferroni's post hoc test (n=3).

**(h, i)** Heat map of reactive transcripts in the (h) Cingulate cortex (CING) and (i) Cerebellum (CBLM) of human DLB and PDD postmortem samples.

**(j-o)** Representative images of double immunostaining for GFAP (green) and C3 (red) in the (j) hippocampus and (m) substantia nigra pars compacta (SNc) in gut injection of α-syn PFF at 2, 6, and 10 month post-injection. Scale bars, 50 μm. Quantification of percentage of $GFAP^+$ cells with DAPI in (k) hippocampus and (n) SNpc and $C3^+$ cells with $GFAP^+$ cells in (l) hippocampus and (o) SNpc. Data are the means ± s.e.m. Two-way ANOVA followed by Bonferroni's post hoc test (n=4).

**(p, q)** Heat map of reactive transcripts in the (p) hippocampus and (q) SNpc of mice injected with PBS or α-syn PFF at 2, 6, 10 months post-injection (n=3). *$P < 0.05$, **$P < 0.01$, ***$P < 0.001$,. n.s., not significant. NTR, neurotoxic reactive astrocytes; NPR, neuroprotective reactive astrocytes.

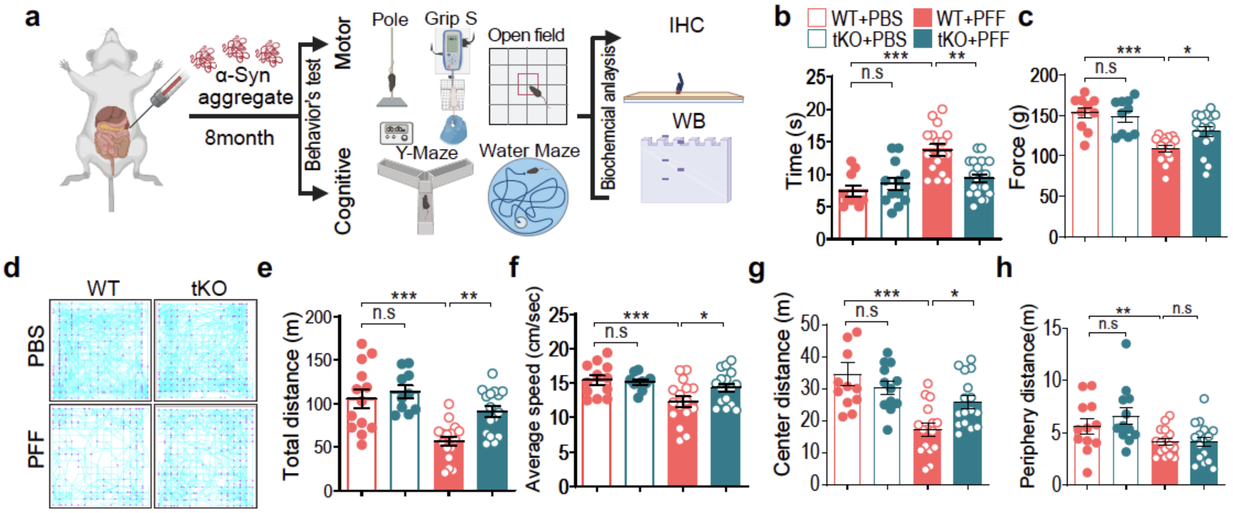


**Extended Data Figure 2. Lewy body dementia-like pathology in the gut-to-brain α-syn PFF model is rescued by genetic depletion of IL1α, TNFα and C1q.**

**(a)** Schematic diagram of the gastrointestinal injection of α-syn PFF or PBS in WT and tKO (*Il1a*, *Tnf* and *C1qa KO*) mice.

**(b-h)** (b) Pole test, (c) grip strength test and (d-h) open field test were performed at 8 months after PBS and α-syn PFF injection. (d) Representative movement paths of mice from each group in the open field test. (e) Total distance, (f) average speed, (g) center distance, and (h) periphery distance were analyzed. Data are the means ± s.e.m. Two-way ANOVA followed by Bonferroni's post hoc test (n=10-16 mice for WT, n=13-16 mice for tKO mice). *P < 0.05, **P < 0.01, ***P < 0.001,. n.s., not significant.

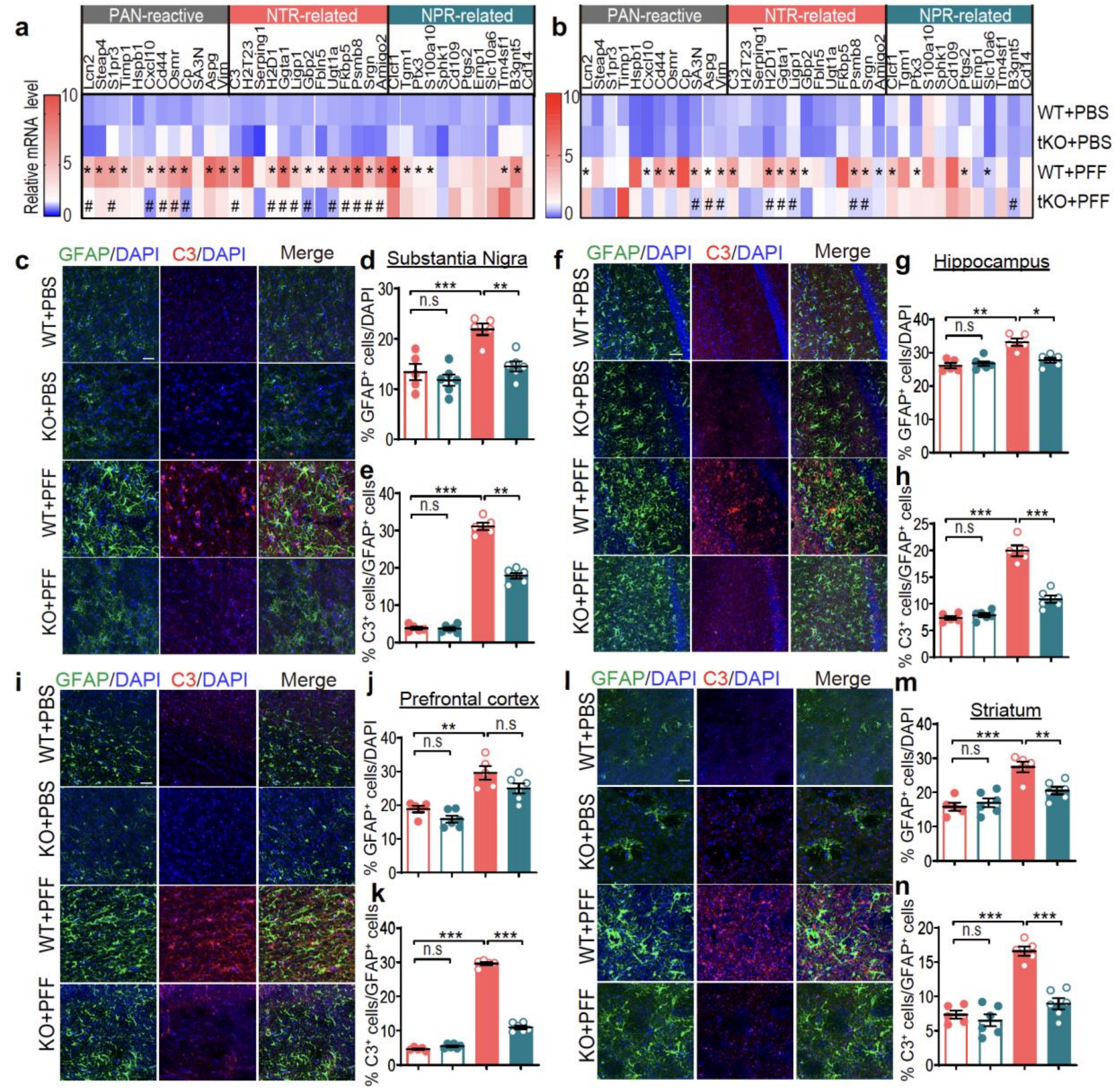


**Extended Data Figure 3. Pathologic $\alpha$-syn-induced NTR astrocyte accumulation is suppressed by genetic depletion of IL1α, TNFα and C1q.**

**(a, b)** Heat map of reactive transcripts in the (a) SNc and (b) hippocampus of WT and tKO (*Il1a*, *Tnf* and *C1qa KO*) mice injected with PBS and α-syn PFF after 10 months.

**(c-n)** Representative images of double immunostaining for GFAP (green) and C3 (red) in the (c) SNc, (f) hippocampus and (i) prefrontal cortex, and (l) striatum. Scale bars, 50 μm. Quantification of percentage of (d, g, j, m) $GFAP^+$ cells with DAPI and (e, h, k, n) $C3^+$ cells with $GFAP^+$ cells in multiple brain regions. Data are the means ± s.e.m. Two-way ANOVA followed by Bonferroni's post hoc test (n=5-6). *$P < 0.05$, **$P < 0.01$, ***$P < 0.001$,. n.s., not significant.

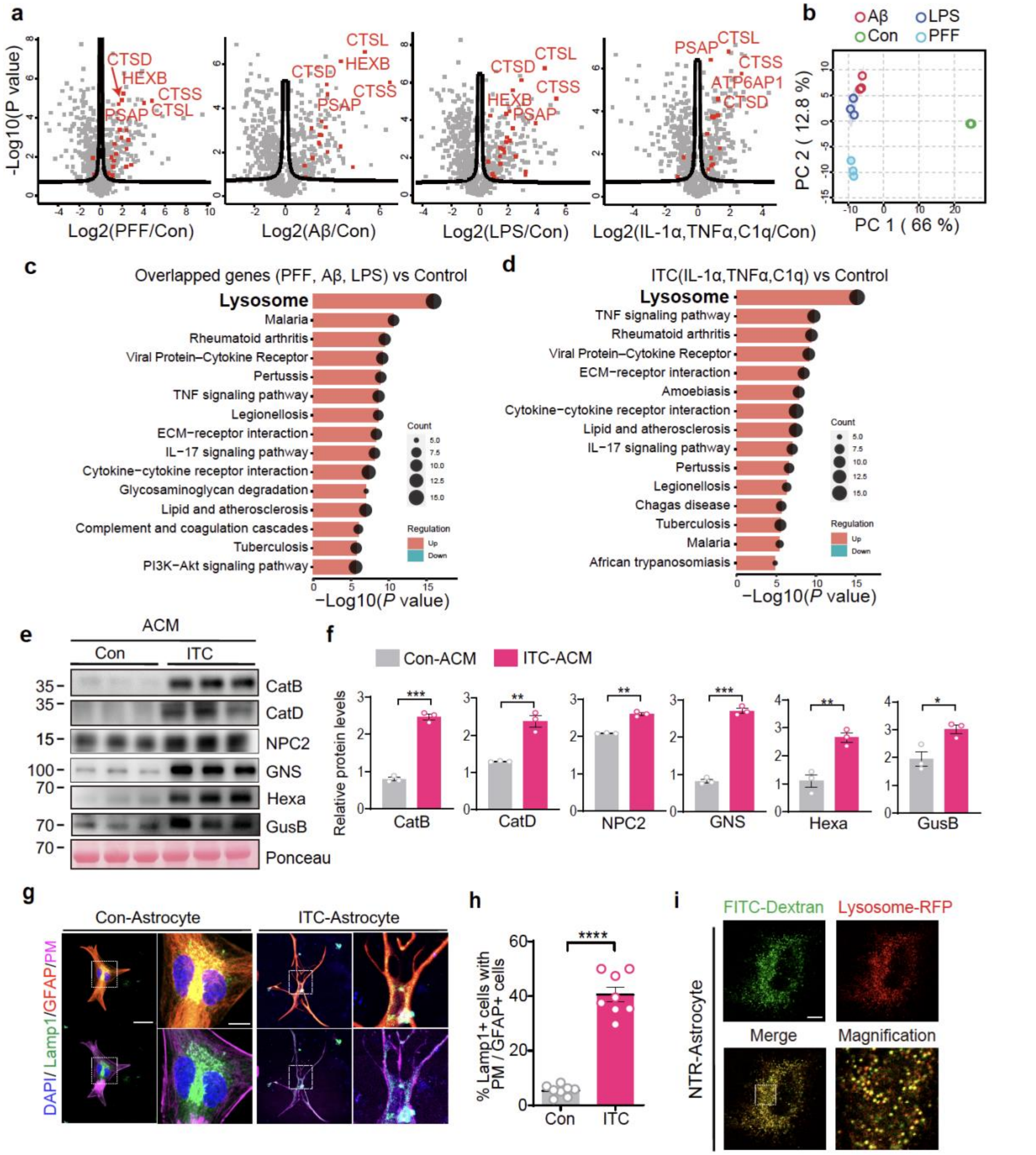


**Extended Data Figure 4. Extracellular release of lysosomal contents from NTR astrocytes.**

**(a)** Changes of secreted proteins from NTR-astrocytes induced by α-syn PFF-activated MCM, Amyloid-β Oligomer-activated MCM, LPS-activated MCM and directly induced by IL-1α, TNFα and C1q cytokine cocktail measured by LC-MS/MS.

**(b)** Graph of Pearson's Correlations of NTR-astrocytes treated with α-syn PFF-MCM, Amyloid-β Oligomer-MCM, LPS-MCM versus control astrocytes.

**(c)** KEGG pathway analyses for the upregulated 102 proteins overlapped in NTR-astrocytes conditioned medium induced by α-syn PFF-MCM, Amyloid-β Oligomer-MCM and LPS-MCM.

**(d)** KEGG pathway analyses between secretome from NTR-astrocyte directly treated with IL-1α , TNFα and C1q and control astrocyte.

**(e, f)** Representative Immunoblots of Cathepsin B and D, NPC2, GNS, GUSB, HEXA and GALNS. Quantification of those proteins level normalized to ponceau S staining. Data are the means ± s.e.m. Unpaired two-tailed student *t* test (n=3).

**(g, h)** Representative images for the positioned lysosome (red, LAMP1) in astrocytes (green, GFAP) treated with IL-1α, TNFα and C1q. Scale bar, 50 μm and 10 μm for low and high magnification images. Quantification of LAMP1 positive cells out of plasma membrane. Data are the means ± s.e.m. Unpaired two-tailed student *t* test (n=8). *P < 0.05, **P < 0.01, ***P < 0.001,. n.s., not significant.

**(i)** Representative images of colocalization of FITC-Dextran and Lysosome-RFP signals in NTR astrocytes. Scale bar, 10 μm.

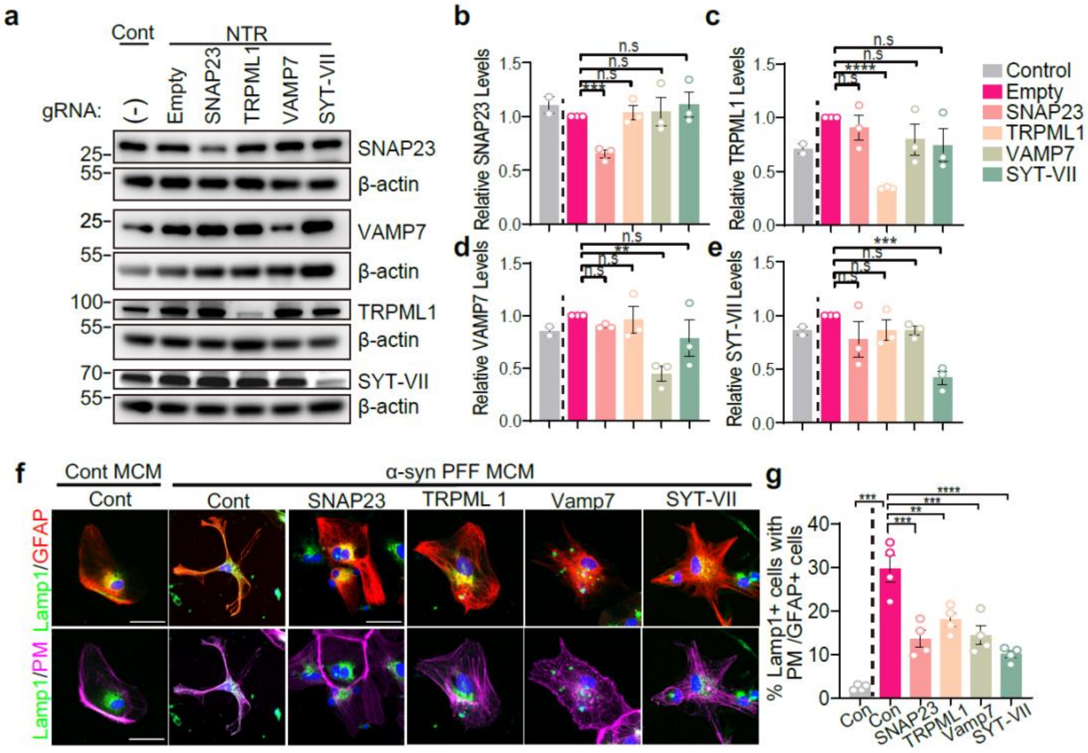


**Extended Data Figure 5. Suppression of lysosomal exocytosis machinery prevents the release lysosomal contents to the extracellular space in astrocytes.**

**(a-g)** Representative immunoblots of SNAP23, TRPML1, VAMP7 and SYT-VII in primary mouse astrocytes infected with lentivirus carrying CRISPR/Cas9 gRNAs targeting mock and lysosomal exocytosis-related genes. (b-e) Quantification of protein levels normalized to β-actin. Data are the means ± s.e.m. Two-way ANOVA followed by Bonferroni's post hoc test (n=3).

**(f, g)** Immunocytochemical analyses for the localization of Lamp1 (green)-positive lysosomes in the plasma membrane (purple) of GFAP-positive astrocytes (red) infected with lentivirus containing CRISPR/Cas9 gRNAs targeting mock and lysosomal exocytosis-related genes. Scale bar, 50 μm. Quantification of Lamp1 positive cells out of plasma membrane. Data are the means ± s.e.m. Two-way ANOVA followed by Bonferroni's post hoc test (n=4). $*P < 0.05$, $**P < 0.01$, $***P < 0.001$,. n.s., not significant.

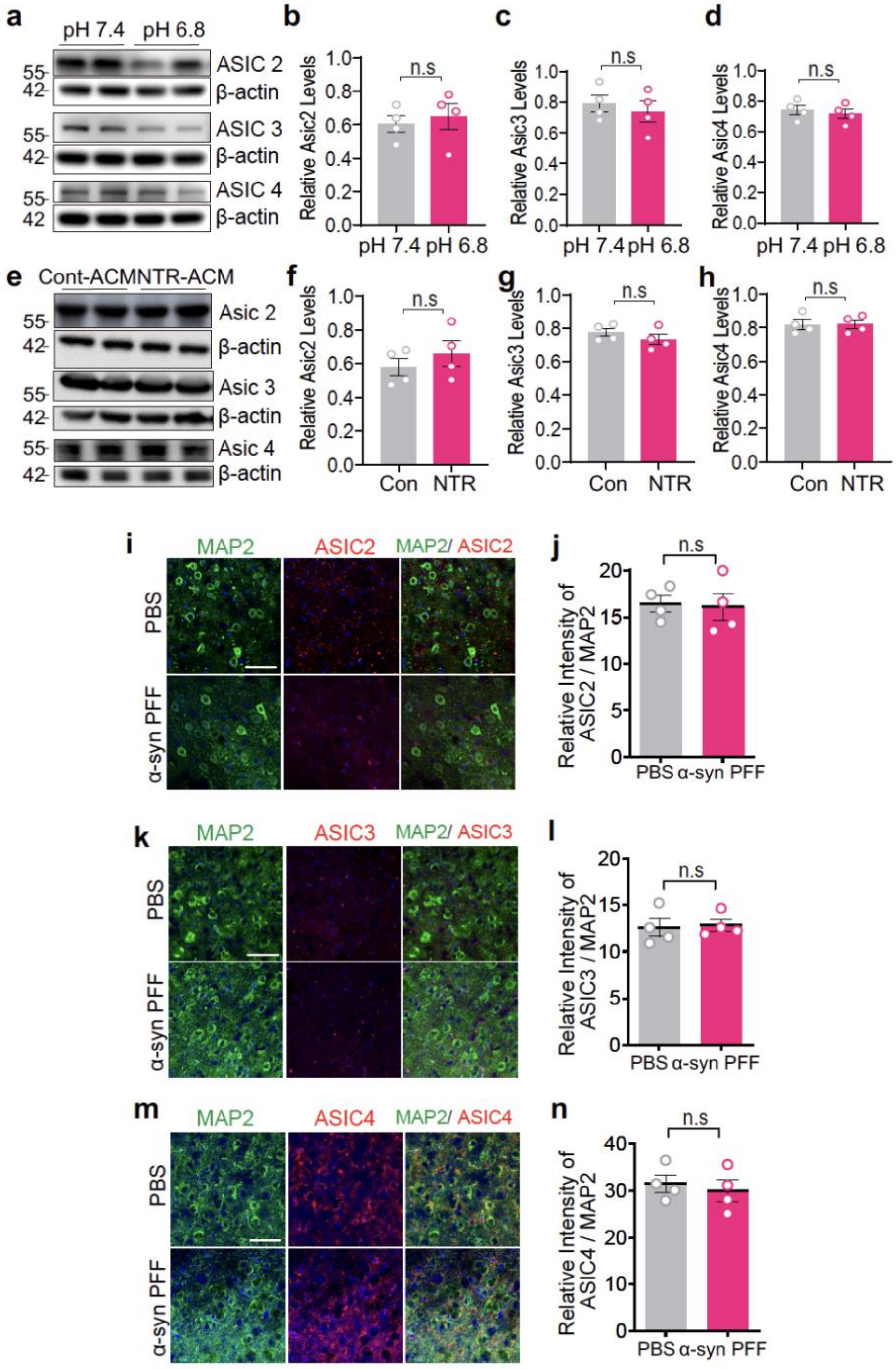


**Extended Data Figure 6. Acidification of extracellular space induced by lysosomal exocytosis by NTR astrocytes does not induce ASIC2-4 expression in neurons.**

**(a-d)** Representative immunoblots of ASIC2,3 and 4 expression in primary mouse cortical neurons incubated with neutral (pH 7.4) or acidic (pH 6.8) medium. Quantification of protein levels normalized β-actin. Data are the means ± s.e.m. Unpaired two-tailed student *t* test (n=4).

**(e-h)** Representative immunoblots of ASIC2,3 and 4 expression in primary mouse cortical neurons treated with control or NTR astrocyte conditioned medium. Quantification of protein levels normalized β-actin. Data are the means ± s.e.m. Unpaired two-tailed student *t* test (n=4). n.s., not significant.

**(i-n)** Representative images of double immunostaining for Map2 (green) and ASIC 2-4 (red) in the SNc, . Scale bar, 50 μm. Quantification of the relative intensity of ASIC $2^{+}$-$4^{+}$ cells within $Map2^{+}$ cells. Data are the means ± s.e.m. Unpaired two-tailed student t test (n=4). n.s., not significant.

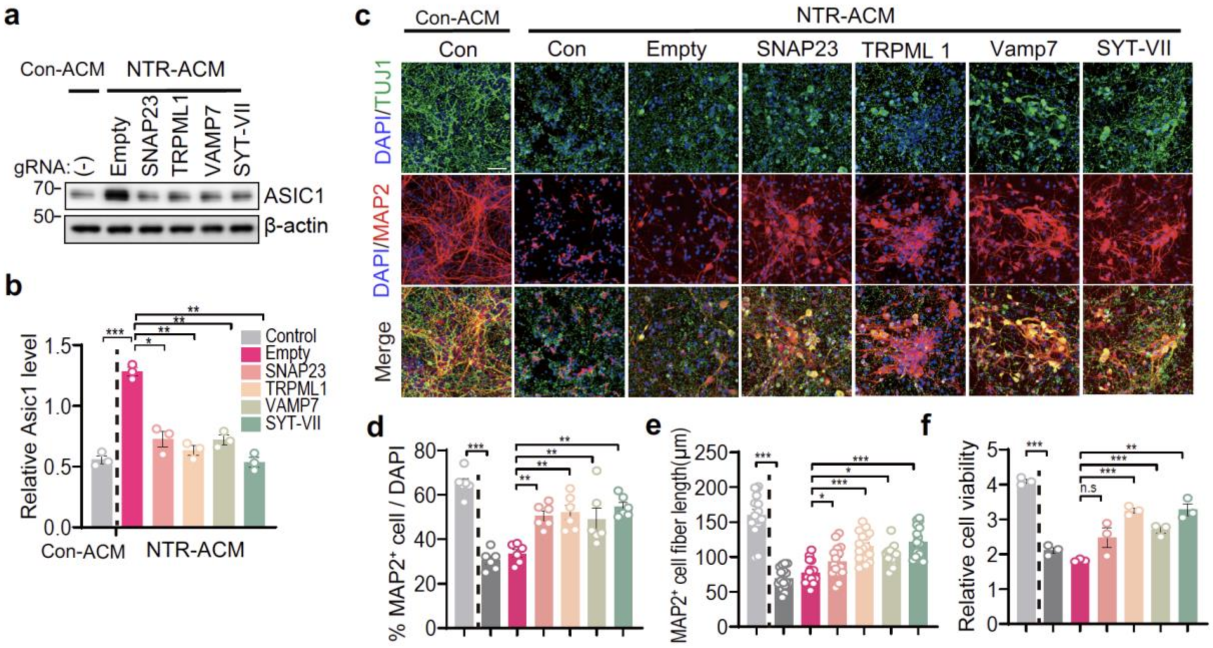


**Extended Data Figure 7. Suppression of lysosomal secretion in NTR astrocyte is sufficient to prevent neuronal ASIC1 expression and relevant neurotoxicity.**

**(a, b)** Representative immunoblots of ASIC1 in primary mouse cortical neurons infected with lentivirus containing CRISPR/Cas9 gRNAs targeting mock and lysosomal exocytosis-related genes followed by incubation with control- or NTR-ACM. Quantification of ASIC1 levels normalized β-actin. Data are the means ± s.e.m. Two-way ANOVA followed by Bonferroni's post hoc test (n=3).

**(c-e)** Representative images of MAP2-positive (red) and TUJ1-positive (green) neurons infected with lentivirus containing CRISPR/Cas9 gRNAs targeting mock and lysosomal exocytosis-related genes followed by incubation with control- or NTR-ACM. Scale bars, 50 μm. (d) percentage of $MAP2^{+}$ cell with DAPI (n=6) and (e) the fiber length (n=16 neurites from 6 independent cultures) were quantified. Data are the means ± s.e.m. Two-way ANOVA followed by Bonferroni's post hoc test.

**(f)** Relative neuronal cell viability was measured using Alamar blue assay. Data are the means ± s.e.m. Two-way ANOVA followed by Bonferroni's post hoc test (n=3). $^{*}P < 0.05$, $^{**}P < 0.01$, $^{***}P < 0.001$,. n.s., not significant.

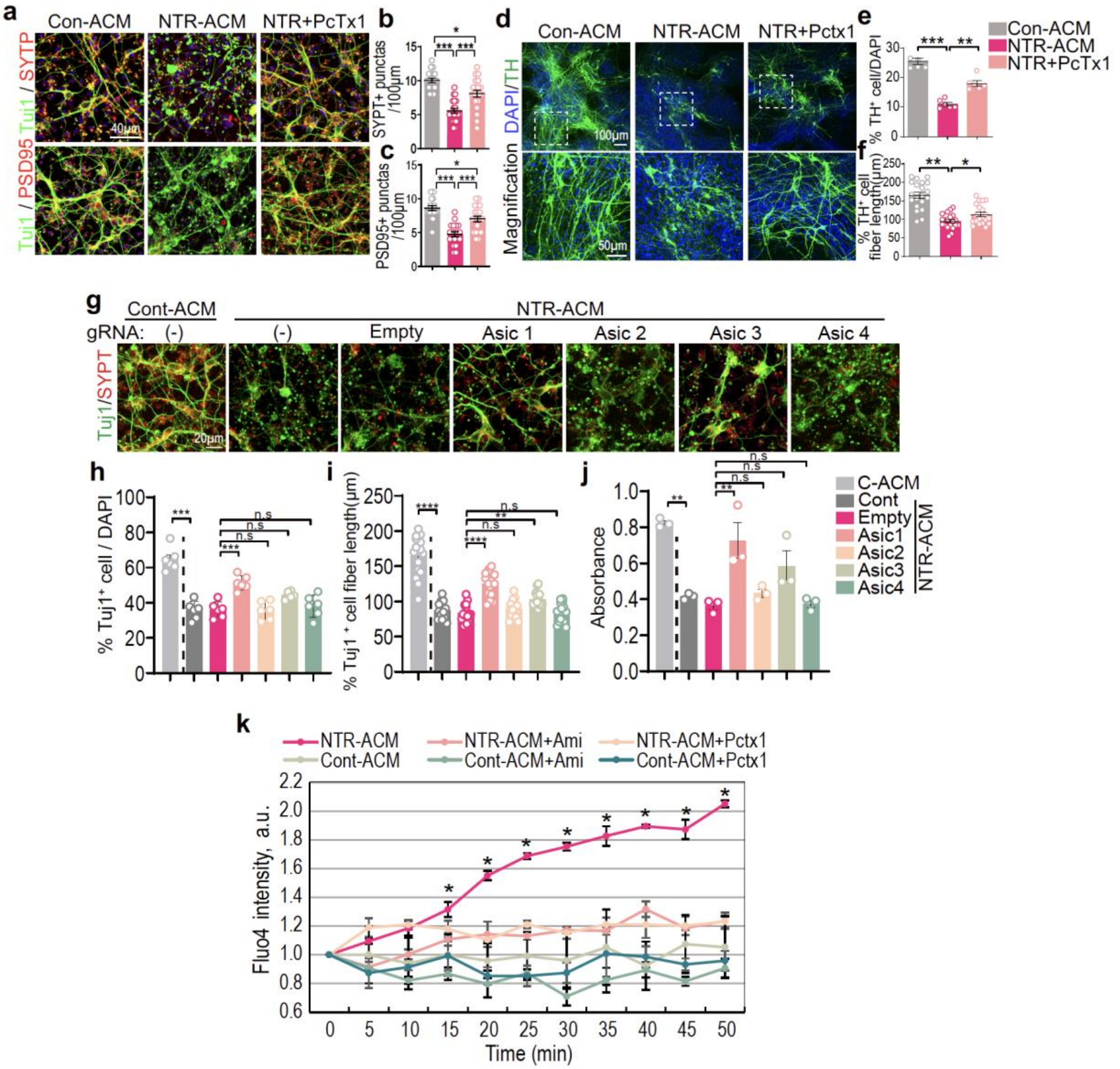


**Extended Data Figure 8. ASIC1 inhibition reduces neuronal cell death, synaptic degeneration, and calcium influx in primary cortical neuron in response to NTR-ACM.**

**(a-c)** Representative immunofluorescence images of presynaptic marker (SYPT) and postsynaptic marker (PSD95) in the differentiated mouse cortical neurons treated with Con-ACM and NTR-ACM with or without Pctx1 or Amiloride. Scale bars, 40 μm. Quantification of (b) Synaptophysin$^{+}$ puncta number in 100 μm TUJ1$^{+}$ fibers and (c) PSD95$^{+}$ puncta number in 100 μm TUJ1$^{+}$ fibers. Data are the means ± s.e.m. Two-way ANOVA followed by Bonferroni's post hoc test (n=23).

**(d-f)** Representative immunofluorescence images of TH (green) and DAPI (blue) in the differentiated primary mouse DA neurons treated with Con-ACM and NTR-ACM with or without Pctx1. Scale bars, 100 μm (top) and 50 μm (bottom). Quantification of (e) TH$^{+}$ cell with DAPI and (f) TH$^{+}$ fiber lengths. Data are the means ± s.e.m. Two-way ANOVA followed by Bonferroni's post hoc test (n=23 from 5 independent cultures).

**(g-i)** Representative images of MAP2-positive (red) and TUJ1-positive (green) cortical neurons infected with lentivirus containing CRISPR/Cas9 gRNAs targeting ASIC1-4 followed by incubation with control- or NTR-ACM. Scale bars, 20 μm. Quantification of (h) TUJ1$^{+}$ cell with DAPI and (i) fiber length. Data are the means ± s.e.m. Two-way ANOVA followed by Bonferroni's post hoc test (n=13-18 from 6 independent cultures).

**(j)** Relative neuronal cell viability was measured using Alamar blue assay. Data are the means ± s.e.m. Two-way ANOVA followed by Bonferroni's post hoc test (n=3).

**(k)** Flouro-4 intensity kinetics graph of cortical neurons measured every 5 minutes until 50 minutes after treatment with Con-ACM or NTR-ACM with vehicle, Pctx1 or Amiloride. Data are the means ± s.e.m. Two-way ANOVA followed by Bonferroni's post hoc test (n=3). $^{*}P < 0.05$, $^{**}P < 0.01$, $^{***}P < 0.001$,. n.s., not significant.

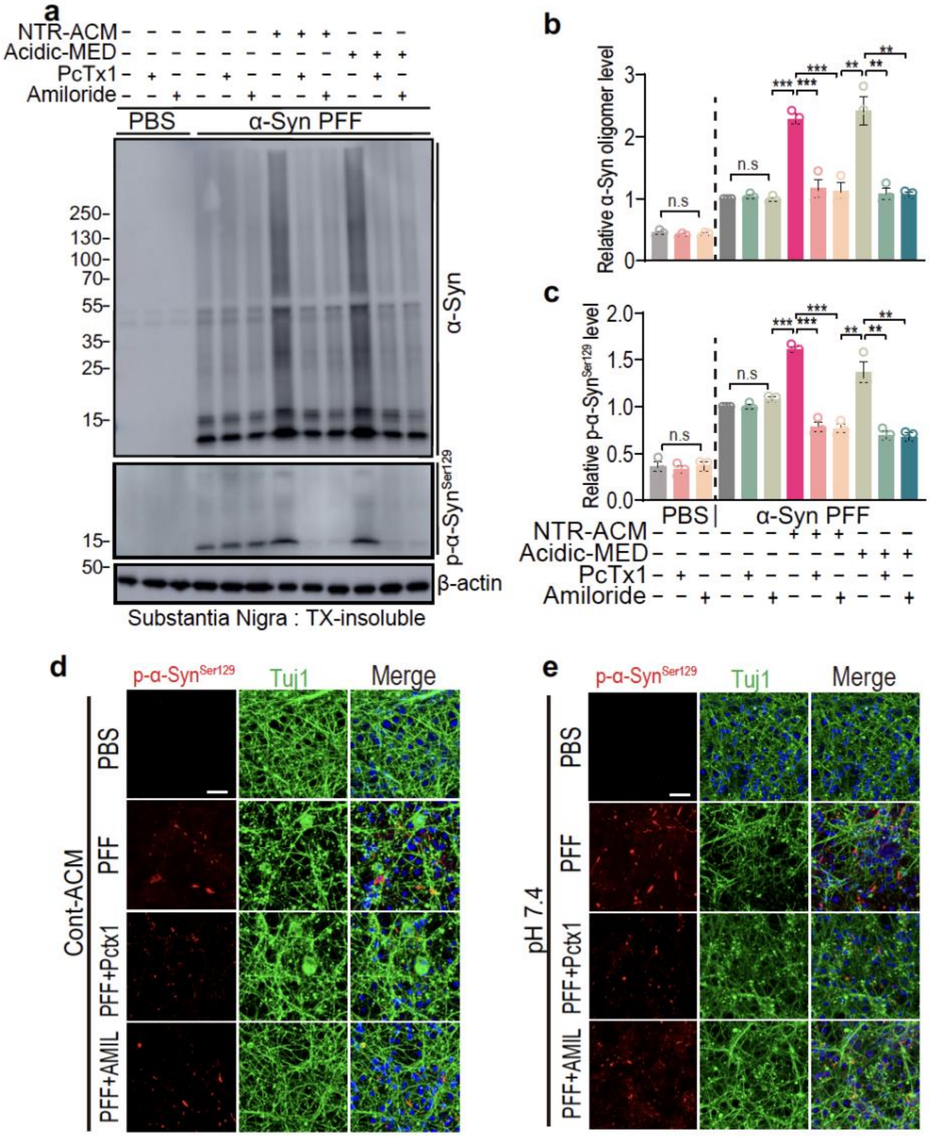


**Extended Data Figure 9. ASIC1 inhibition reduces NTR astrocyte-induced pathologic α-syn accumulation in primary cortical neuron.**

**(a-c)** Representative immunoblots of α-syn, pSer129-α-syn and β-actin in detergent (Triton X-100) insoluble fractions of cortical neurons induced by α-syn PFF treated NTR-ACM or acidic medium (pH 6.8) treated with vehicle or PcTx1 (30 μM for 24h) or Amiloride (100 μM for 24h). Quantification of (b) α-syn oligomer level and (c) pSer129-α-syn level were normalized to β-actin. Data are the means ± s.e.m. Two-way ANOVA followed by Bonferroni's post hoc test (n=3).

**(d, e)** Representative double-immunostaining for pSer129-α-syn (red) and Tuj-1 (green) in primary cortical neurons pre-treated with vehicle, Pctx1 or Amiloride followed by incubation with (d) Con-ACM or (e) neutral medium (pH 7.4). Quantified graph is in Figure 4s and u (n=6). Scale bar, 20 μm.

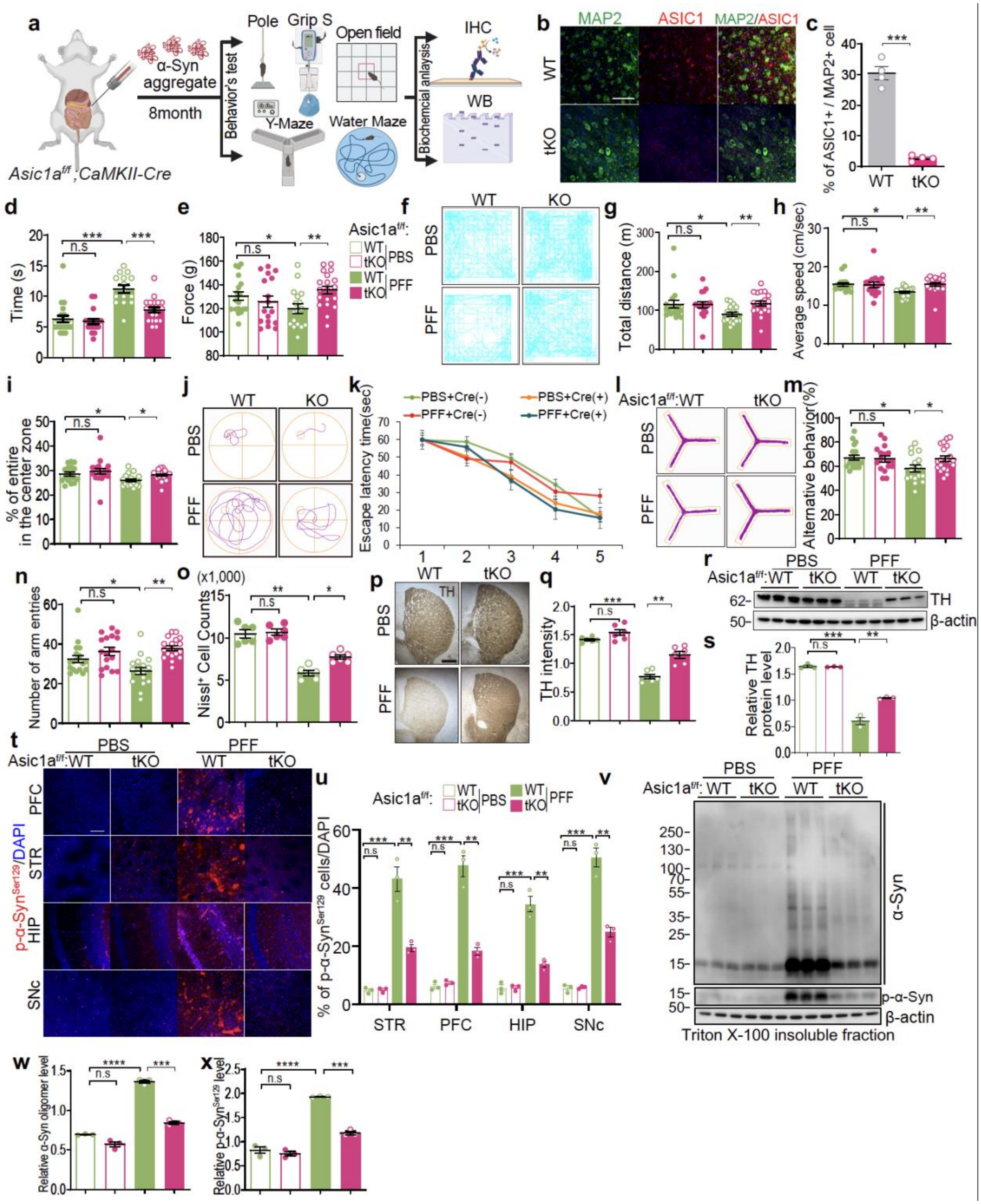

**Extended Data Figure 10. Pathologic α-syn-induced behavioral deficits and neurodegeneration are rescued by genetic depletion of neuronal ASIC1a in vivo.**

**(a)** Schematic diagram of PBS and α-syn PFF gastrointestinal injection in CAMKII-iCre$^-$; ASIC1a$^{flox/flox}$ mice (Control) (n = 17-21), CAMKII-iCre$^+$;ASIC1a$^{flox/flox}$ (Knockout) (n = 18-20).

**(b-c)** Representative images of double immunostaining for Map2 (green) and ASIC1 (red) in the SNc, 50 µm. Quantification of the percentage of ASIC1+ cells within Map2+ cells, n=4.

Results of mice on the **(d)** descend time of pole test **(e)** forelimb force of grip strength test.

**(f-i)** Representative movement paths of mice from each group in the open field test (OFT)., (g) Total distance., (h) average speed (cm/sec)., (i) % of entire in the center zone in open field test.,

**(j-k)** Representative swimming paths of mice from each group in the Morris water maze test (MWMT) on the hidden platform day 5, (k) Result of mice on the Escape latency time in the Morris water maze test (MWMT).

**(l-n)** Representative movement paths of mice from each group in the Y-maze test., (m) Percentage of alternative behavior in the Y-maze test., (n) Number of arm entries in the Y-maze test., **(o)** Nissl positive in the SNpc region.

All behavior tests were analyzed by two-way ANOVA followed by post hoc Bonferroni test for multiple group comparison. *P < 0.05, **P < 0.01, ***P < 0.001,. n.s., not significant.

**(p-q)**. (p) Representative photomicrographs from striatum region containing Vehicle and Amiloride treated mice at 7 months after gut injection of α-syn PFF or PBS. n = 6 biologically independent animals. Scale bar, 1 mm . e,f, quantification graphs of (q) TH-intensity. Data are mean ± s.e.m.; n = 6, biologically independent animals.

**(r-s)** Representative immunoblots of TH normalized to β-actin and Quantification graph.

**(t-u)** Brain distribution of pSer129-α-syn accumulation in mice that received α-syn PFF in the gut. pSer129-α-syn immunohistochemistry from the dorsal motor nucleus of the vagus to the olfactory bulb (Ctx, cortex; DMV, dorsal motor nucleus of the vagus; HIP, hippocampus; LC, locus coeruleus; PFC, prefrontal cortex; SNc, substantia nigra pars compacta; STR, striatum). Data are mean ± s.e.m.; n = 6, biologically independent animals. Scale bar, 100 μm. Quantification of pSer129-α-syn immunoreactivity shown in  (n = 6).

**(v-x)** Representative immunoblots of α-syn andpSer129-α-syn, β-actin in detergent (Triton X-100) insoluble fractions of SNpc. Quantification of (w) α-syn oligomer and (x) pSer129-α-syn protein levels in detergent insoluble fractions. β-actin was used as a loading control.

Data are mean ± s.e.m.; n = 6 biologically independent animals. All biochemical experiments were analyzed by two-way ANOVA followed by post hoc Bonferroni test for multiple group comparison. *P < 0.05, **P < 0.01, ***P < 0.001,. n.s., not significant.

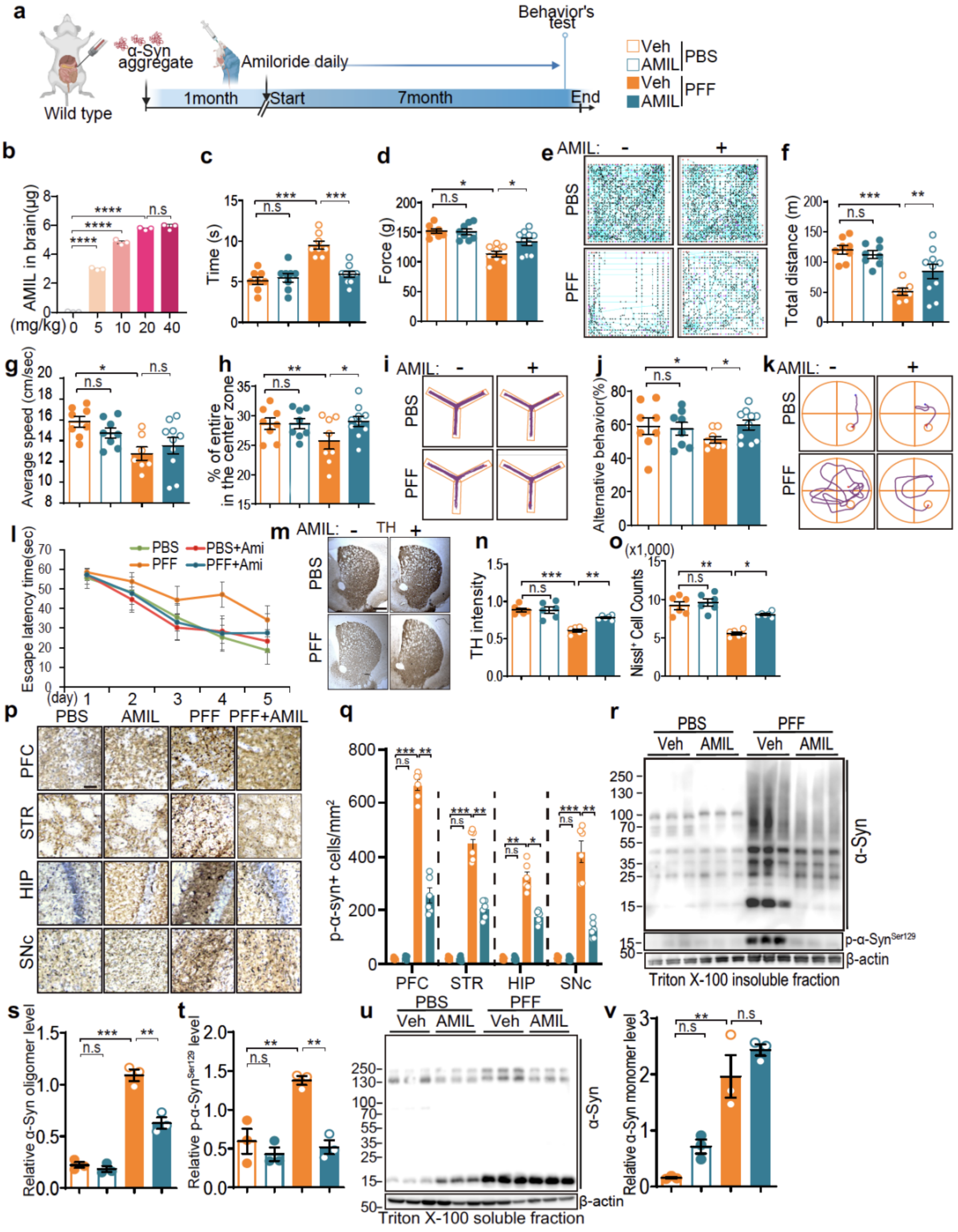

**Extended Data Figure 11. Pathologic α-syn-induced behavioral deficits and neurodegeneration are rescued by pharmacological inhibition of ASIC1 in vivo.**

**(a)** Schematic diagram of Amiloride oral injection experimental design in gut-to-brain α -syn PFF model 7 months after gut injection of α-syn PFF between mice treated with vehicle (PBS) and mice receiving daily oral injections of 5-10 mg $kg^{-1}$ Amiloride (n = 10-12). Created with BioRender.com.

**(b)** Graph of Amiloride level in the brain transferred from Intraperitoneal (IP) injection using HPLC at different concentrations (0, 5, 10, 20, 40 mg/kg of amiloride). Data are the means ± s.e.m. Two-way ANOVA followed by Bonferroni's post hoc test (n=3).

Results of mice on the **(c)** descend time of pole test **(d)** forelimb force of grip strength test.

**(e-h)** Representative movement paths of mice from each group in the open field test(OFT)., (f) Total distance(m) (g) average speed(cm/sec) (h) % of entire in the center zone in open field test.,

**(i-j)** Representative movement paths of mice from each group in the Y-maze test., (j) Percentage of alternative behavior in the Y-maze test.

**(k-l)** Representative swimming paths of mice from each group in the Morris water maze test (MWMT) on the hidden platform day 5.., **(l)** Result of mice on the Escape latency time in the Morris water maze test (MWMT).

**(m-o)** Representative photomicrographs from striatum region containing Vehicle and Amiloride treated mice at 7 months after gut injection of α-syn PFF or PBS. n = 6 biologically independent animals. Scale bar, 1mm. e,f, quantification graphs of (n) TH-intensity in the striatum region. (o) $Nissl^+$ cells positive in the SNpc region. Data are mean ± s.e.m.; n = 6, biologically independent animals.

**(p-q)** Brain distribution of pSer129-α-syn accumulation in mice that received α-syn PFF in the gut. (p) pSer129-α-syn immunohistochemistry from the dorsal motor nucleus of the vagus to the olfactory bulb (Ctx, cortex; DMV, dorsal motor nucleus of the vagus; HIP, hippocampus; LC, locus coeruleus; PFC, prefrontal cortex; SNc, substantia nigra pars compacta; STR, striatum) of Vehicle and Amiloride treated mice at 6 months after gut injection of α-syn PFF or PBS. Data are mean ± s.e.m.; n = 6, biologically independent animals. Scale bar, 20 μm.

(q) Quantification of pSer129-α-syn immunoreactivity shown in (n = 6).

**(r-v)** Representative immunoblots of α-syn, pSer129-α-syn and β-actin in detergent (Triton X-100) (r) insoluble fractions and detergent (u) soluble fractions of SNpc in Vehicle and Amiloride treated mice at 6 months after gut injection of α-syn PFF or PBS. Data are mean ± s.e.m.; n = 3, biologically independent animals., Quantification of (s) α-syn oligomer protein levels, (t) pSer129-α-syn in insoluble and (v) α-syn monomer protein levels, in soluble fractions normalized to β -actin. Data are mean ± s.e.m.; n = 3 biologically independent animals. All biochemical experiments were analyzed by two-way ANOVA followed by post hoc Bonferroni test for multiple group comparison. *P < 0.05, **P < 0.01, ***P < 0.001,. n.s., not significant.

**Table S1. Human post-mortem tissues**

**FFPE tissue sections**

| Group | Diagnosis | Age | Sex | Race | PMD |
|---|---|---|---|---|---|
| Control | 1. Control | 88 | M | W | NA |
| | 2. Control | 88 | F | W | 9 |
| | 3. Control | 69 | F | W | 23 |
| PDD | 1. PD w/Dementia | 71 | M | W | 24 |
| | 2. PD w/Dementia | 73 | F | W | 48 |
| | 3. PD w/Dementia | 75 | M | W | 6 |
| DLB | 1. DLB, mixed Dementia (AD+PD) | 69 | M | W | 24 |
| | 2. DLB, mixed Dementia (AD+PD) | 66 | M | W | 19 |
| | 3. DLB, mixed Dementia (AD+PD) | 74 | M | W | 12 |

**Unfixed frozen tissue**

| Group | Diagnosis | Age | Sex | Race | PMD |
|---|---|---|---|---|---|
| Control | 1. Control | 88 | M | W | NA |
| PDD | 1. PD w/Dementia | 71 | M | W | 24 |
| DLB | 1. DLB, mixed Dementia (AD+PD) | 69 | M | W | 24 |

Abbreviations: PD, Parkinson's disease; W, white; PMD, post-mortem delay (days).

**Table S2. Overlapping proteins identified in NTR astrocyte secretome.**

| PFF | Aβ | LPS | PFF / Aβ | PFF / LPS | Aβ / LPS | PFF/Aβ/LPS |
|---|---|---|---|---|---|---|
| Snca | Lbp | Il1rn | Ctsz | Srgn | Mfap4 | Ccl2 |
| Tnf | Tinagl1 | Pros1 | Ltbp2 | Timp1 | Rarres1 | Cxcl1 |
| Plxnb2 | Cant1 | Gla | Gas1 | Lamp1 | Thbs2 | Sdc4 |
| C1qb | Cx3cl1 | Hist1h4a | Masp1 | Nectin2 | Creg1 | Ccl7 |
| Vcan | Loxl1 | Capg | Cfh | Igfbp5 | C1s1 | Il12b |
| Col4a6 | Mrc1 | Dpp7 | Tf | H2-K1 | Ifi30 | Ctss |
| Itm2b | Aebp1 | Ggh | Col1a2 | Igfbp3 | Atp6ap1 | Csf1 |
| Axl | Plod1 | Manba | Angptl4 | Il1rl1 | Htra1 | Icam1 |
| Cd248 | Col5a1 | Prdx6 | Col5a2 | Il18bp | Col6a1 | Ctsl |
| Smoc1 | | Arhgdia | Loxl2 | Lcp1 | B4galt5 | B2m |
| Tfpi | | Pltp | Col3a1 | Sod1 | Hspg2 | Grn |
| Ppib | | Psma4 | Nid1 | Vnn3 | Fn1 | Saa3 |
| Emilin2 | | Akr1a1 | Col4a2 | Efhd2 | Mmp2 | Spp1 |
| Nhlrc3 | | Pgam1 | | Scg3 | Col12a1 | Ifnar2 |
| Srsf1 | | Plbd2 | | Pik3ip1 | Lamc1 | Apod |
| Try5 | | Gpc6 | | Prdx1 | Lamb1 | Cd5l |
| Mrc2 | | Hist2h3c1 | | Islr | Lama4 | Sirpa |
| Alcam | | Gpi | | Taldo1 | Qsox1 | Procr |
| | | | | | Lgals1 | Vcam1 |
| | | | | | Boc | Serpine1 |
| | | | | | Oaf | Mmp13 |
| | | | | | Col6a3 | Lcn2 |
| | | | | | Tgfbi | Havcr2 |
| | | | | | Lta4h | Cxcl2 |
| | | | | | Smpdl3a | Rnaset2a |
| | | | | | C1qtnf5 | Mmp3 |

| | | | | | | |
|---|---|---|---|---|---|---|
| | | | | | | Sdc1 |
| | | | | | | Fstl1 |
| | | | | | | Ptn |
| | | | | | | H2-Q4 |
| | | | | | | Cp |
| | | | | | | Cd14 |
| | | | | | | Lyz2 |
| | | | | | | Npc2 |
| | | | | | | Gns |
| | | | | | | Lgals3bp |
| | | | | | | Gpnmb |
| | | | | | | Ldlr |
| | | | | | | Ctsb |
| | | | | | | Col18a1 |
| | | | | | | Ftl1 |
| | | | | | | Mxra8 |
| | | | | | | Dcn |
| | | | | | | Tnc |
| | | | | | | Hexb |
| | | | | | | C1ra |
| | | | | | | Il6 |
| | | | | | | Ctsd |
| | | | | | | Galns |
| | | | | | | Cxcl5 |
| | | | | | | Hexa |
| | | | | | | Gm20547 |
| | | | | | | Igfbp7 |
| | | | | | | Psap |

| | | | | | | |
|---|---|---|---|---|---|---|
| | | | | | | Csf1r |
| | | | | | | Cst3 |
| | | | | | | Lum |
| | | | | | | C1qc |
| | | | | | | Ptx3 |
| | | | | | | Nrp2 |
| | | | | | | Fth1 |
| | | | | | | Mertk |
| | | | | | | Man2b1 |
| | | | | | | Thbs1 |
| | | | | | | Gsr |
| | | | | | | Cycs |
| | | | | | | Ccl8 |
| | | | | | | Gm2a |
| | | | | | | Serping1 |
| | | | | | | Col6a2 |
| | | | | | | Gusb |
| | | | | | | Timp2 |
| | | | | | | Ctsc |
| | | | | | | Ppia |
| | | | | | | Clu |
| | | | | | | Ecm1 |
| | | | | | | C3 |
| | | | | | | Lgmn |
| | | | | | | Pdgfrl |
| | | | | | | Rarres2 |
| | | | | | | Loxl3 |
| | | | | | | Pepd |

| | | | | | | | |
|---|---|---|---|---|---|---|---|
| | | | | | | | Uap1l1 |
| | | | | | | | Btd |
| | | | | | | | Bgn |
| | | | | | | | Epdr1 |
| | | | | | | | Tgfbr3 |
| | | | | | | | Plod3 |
| | | | | | | | Chi3l1 |
| | | | | | | | Cpq |
| | | | | | | | Lrp1 |
| | | | | | | | Fbn1 |
| | | | | | | | Pla2g7 |
| | | | | | | | Il4r |
| | | | | | | | Clstn1 |
| | | | | | | | Apoe |
| | | | | | | | Clic1 |
| | | | | | | | Pgk1 |
| | | | | | | | Agrn |
| | | | | | | | Emilin1 |
| | | | | | | | Scpep1 |
| | | | | | | | Igf2 |

**Table S3. Relevant clinical information of control and PD CSF used in Fig. 3j**

| Group | Age | Sex | Group | Age | Sex |
|---|---|---|---|---|---|
| Control (n=29) | 64 | F | PDD/DLB (n=29) | 68 | M |
| | 69 | F | | 73 | M |
| | 52 | M | | 76 | M |
| | 62 | F | | 62 | M |
| | 64 | F | | 70 | M |
| | 72 | F | | 65 | M |
| | 63 | M | | 80 | F |
| | 57 | M | | 78 | F |
| | 75 | M | | 67 | F |
| | 75 | M | | 69 | F |
| | 67 | M | | 71 | M |
| | 60 | M | | 71 | F |
| | 72 | M | | 48 | F |
| | 60 | M | | 65 | M |
| | 59 | M | | 60 | F |
| | 70 | F | | 71 | M |
| | 67 | F | | 74 | M |
| | 65 | M | | 66 | M |
| | 49 | M | | 68 | M |
| | 62 | M | | 84 | M |
| | 55 | F | | 53 | M |
| | 67 | F | | 68 | F |
| | 56 | F | | 75 | M |
| | 57 | M | | 48 | M |
| | 55 | F | | 77 | F |

| | | | | | |
|---|---|---|---|---|---|
| | 65 | M | | 66 | M |
| | 53 | M | | 74 | M |
| | 62 | F | | 43 | F |
| | 57 | F | | 75 | M |
| **Summarized information** | | | | | |
| **Group** | **Mean age ± s.e.m.** | **F/M ratio (%)** | **Group** | **Mean age ± s.e.m.** | **F/M ratio (%)** |
| Control | 62.1 ± 1.24 | 48.3 / 51.7 | PD | 67.0 ± 1.69 | 31 / 69 |

**Table S4. Reagent or resources used in this study**

| REAGENT or RESOURCE | SOURCE | IDENTIFIER |
|---|---|---|
| Mouse monoclonal Anti-glial fibrillary acidic protein (GFAP) antibody | MP Biomedicals | Cat# 0869110-CF |
| Rabbit polyclonal Anti-Glial Fibrillary Acidic Protein | DAKO | Cat# Z033429-2 |
| Mouse monoclonal anti-C3 Antibody (B-9) | Santa Cruz Biotechnology | Cat# sc-28294 |
| Mouse monoclonal anti-cathepsin B Antibody (H-5) | Santa Cruz Biotechnology | Cat# sc-365558 |
| Mouse monoclonal anti-cathepsin D Antibody (C-5) | Santa Cruz Biotechnology | Cat# sc-377124 |
| Rabbit polyclonal anti-Npc2 Polyclonal Antibody | Thermo Fisher Scientific | Cat# PA5-51463 |
| Rabbit polyclonal Anti-GNS Antibody | Proteintech | Cat# 13044-1-AP |
| Rabbit polyclonal Anti-GUSB Antibody | Proteintech | Cat# 16332-1-AP |
| Rabbit polyclonal anti-GALNS Antibody | ThermoFisher Scientific | Cat# PA522098 |
| Rabbit polyclonal anti-HEXA Antibody | ThermoFisher Scientific | Cat# PA5102823 |
| Mouse monoclonal anti-b-actin-peroxidase (clone AC-15) antibody | Sigma | Cat# A3854 |
| Rabbit polyclonal Anti-LAMP1 antibody | Abcam | Cat# ab24170 |
| Rabbit polyclonal Anti-MAP2 Antibody | Proteintech | Cat# 17490-1-AP |
| Mouse monoclonal anti-a-synuclein antibody | BD Bioscience | Cat# 610787 |
| Mouse monoclonal anti-a-synuclein phospho(Ser129) antibody | Biolegend | Cat# 825701 |
| Rabbit polyclonal anti-Tyrosine Hydroxylase antibody | Novus Biologicals | Cat# NB300-109 |
| Mouse monoclonal anti-SLC6A3/DAT1 antibody (mAb16) | Novus Biologicals | Cat# NBP2-22164SS |

| | | |
|---|---|---|
| Rabbit polyclonal anti-Tubulin β-3 (TUBB3) Antibody | Biolegend | Cat# 801201 |
| Mouse monoclonal anti-Tubulin β-3 (TUBB3) Antibody | Biolegend | Cat# 802001 |
| Rabbit polyclonal anti-ASIC1 Antibody | ThermoFisher Scientific | Cat# PA5-26278 |
| Mouse monoclonal anti- ASIC 1 Antibody | Biolegend | Cat# 833501 |
| Rabbit polyclonal anti- ASIC 2 Antibody | ThermoFisher Scientific | Cat# OSR00098W |
| Rabbit polyclonal anti- ASIC 3 Antibody | ThermoFisher Scientific | Cat# PA5-77734 |
| Rabbit polyclonal anti- ASIC 4 Antibody | ThermoFisher Scientific | Cat# OSR00101W |
| Mouse monoclonal anti-Synaptophysin antibody | BD Biosciences | Cat# 611880 |
| Rabbit polyclonal anti-PSD95 antibody | Abcam | Cat# ab18258 |
| Goat anti-Rabbit IgG (H+L) Secondary antibody Alexa Fluor 488 | ThermoFisher Scientific | Cat# A11008 |
| Goat anti-Rabbit IgG (H+L) Secondary antibody Alexa Fluor 594 | ThermoFisher Scientific | Cat# A11037 |
| Goat anti-mouse IgG (H+L) Secondary antibody Alexa Fluor 488 antibody | ThermoFisher Scientific | Cat# A11029 |
| Goat anti-mouse IgG (H+L) Secondary antibody Alexa Fluor 594 antibody | ThermoFisher Scientific | Cat# A11005 |
| Cellbrite Fix 640 CellBrite Fix Membrane stains | Biotium | Cat# 30089-T |
| FITC-dextran | Chondrex | Cat# CFD150 |
| Molecular Probes CellLight Lysosomes RFP | ThermoFisher Scientific | Cat# C10597 |

**Table S5. Primers used in this study**

**Mouse primer sequences.**

| Genes | Forward primer | Reverse primer | Size (bp) |
|---|---|---|---|
| ASIC1 | CTCCGGAGCAGTACAAGGAG | GTACTTGGCTGAGGCTTTGC | 155 |
| ASIC2 | GAGGCGCTCAATTACGAGAC | ATCATGGCTCCCTTCCTCTT | 201 |
| ASIC3 | TGAGAGCCACCAGCTTACCT | ACATGTCCTCAAGGGAGTGG | 245 |
| ASIC4 | ATGGTCAAGATCCCCAACAG | AATGAACAGGCCCATCTGTC | 198 |
| Lcn2 | CCAGTTCGCCATGGTATTTT | CACACTCACCACCCATTCAG | 206 |
| Steap4 | CCCGAATCGTGTCTTTCCTA | GGCCTGAGTAATGGTTGCAT | 262 |
| S1pr3 | AAGCCTAGCGGGAGAGAAAC | TCAGGGAACAATTGGGAGAG | 197 |
| Timp1 | AGTGATTTCCCCGCCAACTC | GGGGCCATCATGGTATCTGC | 2088 |
| Hspb1 | GACATGAGCAGTCGGATTGA | GGATGGGGTGTAGGGGTACT | 265 |
| Cxcl10 | CCCACGTGTTGAGATCATTG | CACTGGGTAAAGGGGAGTGA | 211 |
| Cd44 | ACCTTGGCCACCACTCCTAA | GCAGTAGGCTGAAGGGTTGT | 299 |
| Osmr | GTGAAGGACCCAAAGCATGT | GCCTAATACCTGGTGCGTGT | 199 |
| Cp | TGTGATGGGAATGGGCAATGA | AGTGTATAGAGGATGTTCCAGGTCA | 282 |
| Serpinga3n | CCTGGAGGATGTCCTTTCAA | TTATCAGGAAAGGCCGATTG | 233 |
| Aspg | GCTGCTGGCCATTTACACTG | GTGGGCCTGTGCATACTCTT | 133 |
| Vim | AGACCAGAGATGGACAGGTGA | TTGCGCTCCTGAAAAACTGC | 169 |
| Gfap | AGAAAGGTTGAATCGCTGGA | CGGCGATAGTCGTTAGCTTC | 299 |
| C3 | CCAGCTCCCCATTAGCTCTG | GCACTTGCCTCTTTAGGAAGTC | 159 |
| H2-T23 | GGACCGCGAATGACATAGC | GCACCTCAGGGTGACTTCAT | 212 |
| Serping1 | ACAGCCCCCTCTGAATTCTT | GGATGCTCTCCAAGTTGCTC | 299 |
| H2-D1 | TCCGAGATTGTAAAGCGTGAAGA | ACAGGGCAGTGCAGGGATAG | 204 |
| Ggta1 | GTGAACAGCATGAGGGGTTT | GTTTTGTTGCCTCTGGGTGT | 115 |
| Ligp1 | GGGGCAATAGCTCATTGGTA | ACCTCGAAGACATCCCCTTT | 102 |
| Gbp2 | GGGGTCACTGTCTGACCACT | GGGAAACCTGGGATGAGATT | 285 |

| | | | |
|---|---|---|---|
| Fbln5 | CTTCAGATGCAAGCAACAA | AGGCAGTGTCAGAGGCCTTA | 281 |
| Ugt1a | CCTATGGGTCACTTGCCACT | AAAACCATGTTGGGCATGAT | 136 |
| Fkbp5 | TATGCTTATGGCTCGGCTGG | CAGCCTTCCAGGTGGACTTT | 194 |
| Psmb8 | CAGTCCTGAAGAGGCCTACG | CACTTTCACCCAACCGTCTT | 121 |
| Srgn | GCAAGGTTATCCTGCTCGGA | TGGGAGGGCCGATGTTATTG | 134 |
| Amigo2 | GAGGCGACCATAATGTCGTT | GCATCCAACAGTCCGATTCT | 263 |
| Clcf1 | CTTCAATCCTCCTCGACTGG | TACGTCGGAGTTCAGCTGTG | 176 |
| Tgm1 | CTGTTGGTCCCGTCCCAAA | GGACCTTCCATTGTGCCTGG | 97 |
| Ptx3 | AACAAGCTCTGTTGCCCATT | TCCCAAATGGAACATTGGAT | 147 |
| S100a10 | CCTCTGGCTGTGGACAAAAT | CTGCTCACAAGAAGCAGTGG | 238 |
| Sphk1 | GATGCATGAGGTGGTGAATG | TGCTCGTACCCAGCATAGTG | 135 |
| Cd109 | CACAGTCGGGAGCCCTAAAG | GCAGCGATTTCGATGTCCAC | 147 |
| Ptgs2 | GCTGTACAAGCAGTGGCAAA | CCCCAAAGATAGCATCTGGA | 232 |
| Emp1 | GAGACACTGGCCAGAAAAGC | TAAAAGGCAAGGGAATGCAC | 183 |
| Slc10a6 | GCTTCGGTGGTATGATGCTT | CCACAGGCTTTTCTGGTGAT | 217 |
| Tm4sf1 | GCCCAAGCATATTGTGGAGT | AGGGTAGGATGTGGCACAAG | 258 |
| B3gnt5 | CGTGGGGCAATGAGAACTAT | CCCAGCTGAACTGAAGAAGG | 207 |
| Cd14 | GGACTGATCTCAGCCCTCTG | GCTTCAGCCCAGTGAAAGAC | 232 |

**Human primer sequences.**

| Genes | Forward primer | Reverse primer | Size (bp) |
|---|---|---|---|
| Lcn2 | CCACCTCAGACCTGATCCCA | CCCCTGGAATTGGTTGTCCTG | 80 |
| Steap4 | GGCTTTGGGAATACTTGGGTT | TGGACAAATCGGAACTCTCTCC | 102 |
| S1pr3 | CGGCATCGCTTACAAGGTCAA | GCCACGAACATACTGCCCT | 99 |
| Timp1 | CTTCTGCAATTCCGACCTCGT | ACGCTGGTATAAGGTGGTCTG | 79 |
| Hspb1 | ACGGTCAAGACCAAGGATGG | AGCGTGTATTTCCGCGTGA | 104 |
| Cxcl10 | GTGGCATTCAAGGAGTACCTC | TGATGGCCTTCGATTCTGGATT | 198 |

| | | | |
|---|---|---|---|
| Cd44 | CTGCCGCTTTGCAGGTGTA | CATTGTGGGCAAGGTGCTATT | 109 |
| Osmr | ATGGCTCTATTTGCAGTCTTTCA | CACCCAGATGACATTGGATGTT | 246 |
| Cp | GGGCCATCTACCCTGATAACA | TTAAAGGTCCGATGAGTCCTGA | 198 |
| Serpinga3n | CCTGAAGGCCCCTGATAAGAA | GCTGGACTGATTGAGGGTGC | 196 |
| Aspg | AACCGGGCAACCAAGGTAG | CCAGCTCCCTGTTGATTGTGA | 106 |
| Vim | GACGCCATCAACACCGAGTT | CTTTGTCGTTGGTTAGCTGGT | 238 |
| Gfap | CTGCGGCTCGATCAACTCA | TCCAGCGACTCAATCTTCCTC | 209 |
| C3 | GGGGAGTCCCATGTACTCTATC | GGAAGTCGTGGACAGTAACAG | 125 |
| H2-T23 | TTCCGAGTGAATCTGCGGAC | GTCGTAGGCGAACTGTTCATAC | 138 |
| Serping1 | CTGGCTGGGGATAGAGCCT | GAGATAACTGTTGTTGCGACCT | 110 |
| H2-D1 | ACCCTCGTCCTGCTACTCTC | CTGTCTCCTCGTCCCAATACT | 244 |
| Ggta1 | AGAGGAGACCAAAGGAAGGAAA | GGATTAAACCAGTCCCATAGCC | 62 |
| Ligp1 | GAAGGAGGCATCCAATAGCAG | ACTCTCGGACACCACTCCATT | 85 |
| Gbp2 | CTATCTGCAATTACGCAGCCT | TGTTCTGGCTTCTTGGGATGA | 182 |
| Fbln5 | CTCACTGTTACCATTCTGGCTC | GACTGGCGATCCAGGTCAAAG | 89 |
| Ugt1a | CATGCTGGGAAGATACTGTTGAT | GCCCGAGACTAACAAAAGACTCT | 214 |
| Fkbp5 | AATGGTGAGGAAACGCCGATG | TCGAGGGAATTTTAGGGAGACT | 250 |
| Psmb8 | CACGCTCGCCTTCAAGTTC | AGGCACTAATGTAGGACCCAG | 80 |
| Srgn | AGGTTATCCTACGCGGAGAG | GTCTTTGGAAAAAGGTCAGTCCT | 156 |
| Amigo2 | CCTGGGAACCTTTTCAGACTG | GCAAACGATACTGGAATCCACT | 92 |
| Clcf1 | TTTCAACGAGCCAGACTTCAAC | GAGGCCACGCAAGTAACACA | 157 |
| Tgm1 | GCACCACACAGACGAGTATGA | GGTGATGCGATCAGAGGATTC | 109 |
| Ptx3 | CATCTCCTTGCGATTCTGTTTTG | CCATTCCGAGTGCTCCTGA | 165 |
| S100a10 | GGCTACTTAACAAAGGAGGACC | GAGGCCCGCAATTAGGGAAA | 168 |
| Sphk1 | GCTCTGGTGGTCATGTCTGG | CACAGCAATAGCGTGCAGT | 209 |
| Cd109 | AAGCCAGTGAAAGGAGACGTA | CCAGGGGAAGATAGATCCAGG | 185 |
| Ptgs2 | CTGGCGCTCAGCCATACAG | CGCACTTATACTGGTCAAATCCC | 94 |
| Emp1 | GTGCTGGCTGTGCATTCTTG | CCGTGGTGATACTGCGTTCC | 81 |

| | | | |
|---|---|---|---|
| Slc10a6 | GGAAGCTGTGGTCGCACAT | GTAAAAGGCATGAGCCCAAACT | 79 |
| Tm4sf1 | TGCATCGGACATTCTCTGGTG | GTTCCAGCCCAATGAAGACAA | 190 |
| B3gnt5 | GGGCCTCGCTACCAATACTTG | CGGAACGTCGATCATAGTTTTCA | 109 |
| Cd14 | ACGCCAGAACCTTGTGAGC | GCATGGATCTCCACCTCTACTG | 122 |

### Guide RNAs

| Name | Sequence |
|---|---|
| gRNAs targeting ASIC1-#1 forward | CACCGACTGCTCCGGAGTACAGTAT |
| gRNAs targeting ASIC1-#1 reverse | AAACATACTGTACTCCGGAGCAGTC |
| gRNAs targeting ASIC1-#2 forward | CACCGGGTTTCACAATCAATCCGGC |
| gRNAs targeting ASIC1-#2 reverse | AAACGCCGGATTGATTGTGAAACCC |
| gRNAs targeting ASIC1-#3 forward | CACCGCAAACGTGCCTCGAGCGGGA |
| gRNAs targeting ASIC1-#3 reverse | AAACTCCCGCTCGAGGCACGTTTGC |
| gRNAs targeting ASIC2-#1 forward | CACCGGGTCTCACAGTCGATCCGAC |
| gRNAs targeting ASIC2-#1 reverse | AAACGTCGGATCGACTGTGAGACCC |
| gRNAs targeting ASIC2-#2 forward | CACCGATATCTGCCCCCGCCGTGGG |
| gRNAs targeting ASIC2-#2 reverse | AAACCCCACGGCGGGGGCAGATATC |
| gRNAs targeting ASIC2-#3 forward | CACCGTCACATATCTGCCCCCGCCG |
| gRNAs targeting ASIC2-#3 reverse | AAACCGGCGGGGGCAGATATGTGAC |
| gRNAs targeting ASIC3-#1 forward | CACCGGCGGTATTGCAGTCACCCCA |
| gRNAs targeting ASIC3-#1 reverse | AAACTGGGGTGACTGCAATACCGCC |
| gRNAs targeting ASIC3-#2 forward | CACCGTGAGCGGGTTCGCTACTATG |
| gRNAs targeting ASIC3-#2 reverse | AAACCATAGTAGCGAACCCGCTCAC |
| gRNAs targeting ASIC3-#3 forward | CACCGTCGGATCCCCACCTCAAATG |
| gRNAs targeting ASIC3-#3 reverse | AAACCATTTGAGGTGGGGATCCGAC |
| gRNAs targeting ASIC4-#1 forward | CACCGTGCAGACGAGACGTCATTCG |
| gRNAs targeting ASIC4-#1 reverse | AAACCGAATGACGTCTCGTCTGCAC |
| gRNAs targeting ASIC4-#2 forward | CACCGCCGGGCGGAAAGCGAGCTCA |

| | |
|---|---|
| gRNAs targeting ASIC4-#2 reverse | AAACTGAGCTCGCTTTCCGCCCGGC |
| gRNAs targeting ASIC4-#3 forward | CACCGTCTCTTTGCCGTAACGAGTC |
| gRNAs targeting ASIC4-#3 reverse | AAACGACTCGTTACGGCAAAGAGAC |
| gRNAs targeting SYT7-#1 forward | CACCGTTTCGCCTTCGACATACCCA |
| gRNAs targeting SYT7-#1 reverse | AAACTGGGTATGTCGAAGGCGAAAC |
| gRNAs targeting SYT7-#2 forward | CACCGGTGGGGCAGATTCGAAACCG |
| gRNAs targeting SYT7-#2 reverse | AAACCGGTTTCGAATCTGCCCCACC |
| gRNAs targeting SYT7-#3 forward | CACCGACAACGTGTGTGGCGGAGCG |
| gRNAs targeting SYT7-#3 reverse | AAACCGCTCCGCCACACACGTTGTC |
| gRNAs targeting TRPML1-#1 forward | CACCGGTATTCATGTGGCGGCGGCG |
| gRNAs targeting TRPML1-#1 reverse | AAACCGCCGCCGCCACATGAATACC |
| gRNAs targeting TRPML1-#2 forward | CACCGTCTGTGGCTGGATCGTTCTA |
| gRNAs targeting TRPML1-#2 reverse | AAACTAGAACGATCCAGCCACAGAC |
| gRNAs targeting TRPML1-#3 forward | CACCGCAGCAGCGAATGGTCCTCCG |
| gRNAs targeting TRPML1-#3 reverse | AAACCGGAGGACCATTCGCTGCTGC |
| gRNAs targeting VAMP7-#1 forward | CACCGCCACGTTGAGCAACTAAATC |
| gRNAs targeting VAMP7-#1 reverse | AAACGATTTAGTTGCTCAACGTGGC |
| gRNAs targeting VAMP7-#2 forward | CACCGAACTCGCTATTCATAGCATA |
| gRNAs targeting VAMP7-#2 reverse | AAACTATGCTATGAATAGCGAGTTC |
| gRNAs targeting VAMP7-#3 forward | CACCGTAGCGAGTTTTCAAGTGTTT |
| gRNAs targeting VAMP7-#3 reverse | AAACAAACACTTGAAAACTCGCTAC |
| gRNAs targeting SNAP23-#1 forward | CACCGTGACCATTTGTAATCCGGCT |
| gRNAs targeting SNAP23-#1 reverse | AAACAGCCGGATTACAAATGGTCAC |
| gRNAs targeting SNAP23-#2 forward | CACCGCAACCGAGCCGGATTACAAA |
| gRNAs targeting SNAP23-#2 reverse | AAACTTTGTAATCCGGCTCGGTTGC |
| gRNAs targeting SNAP23-#3 forward | CACCGTAGTATCTAAGCAACCGAGC |
| gRNAs targeting SNAP23-#3 reverse | AAACGCTCGGTTGCTTAGATACTAC |
